# Behavioral timescale synaptic plasticity in the hippocampus creates non-spatial representations during learning and is modulated by entorhinal inputs

**DOI:** 10.1101/2024.08.27.609983

**Authors:** Conor C. Dorian, Jiannis Taxidis, Ahmet Arac, Peyman Golshani

## Abstract

Behavioral timescale synaptic plasticity (BTSP) is a form of synaptic potentiation where a single plateau potential in hippocampal neurons forms a place field during spatial learning. We asked whether BTSP can also form non-spatial responses in the hippocampus and what roles the medial and lateral entorhinal cortex (MEC and LEC) play in driving non-spatial BTSP. Two-photon calcium imaging of dorsal CA1 neurons while mice performed an odor-cued working memory task revealed plateau-like events which formed stable odor-specific responses. These BTSP-like events were much more frequent during the first day of task learning, suggesting that BTSP may be important for early learning. Strong single-neuron stimulation through holographic optogenetics induced plateau-like events and subsequent odor-fields, causally linking BTSP with non-spatial representations. MEC chemogenetic inhibition reduced the frequency of plateau-like events, whereas LEC inhibition reduced potentiation and field-induction probability. Calcium imaging of LEC and MEC temporammonic CA1 projections revealed that MEC axons were more strongly activated by odor presentations, while LEC axons were more odor-selective, further confirming the role of MEC in driving plateau-like events and LEC in relaying odor-specific information. Altogether, odor-specific information from LEC and strong odor-timed activity from MEC are crucial for driving BTSP in CA1, which is a synaptic plasticity mechanism for generation of both spatial and non-spatial responses in the hippocampus.

## INTRODUCTION

In many situations, learning is not a gradual process. In fact, our ability to make associations after a single experience is critical for survival. While there have been dramatic improvements in artificial intelligence and machine learning algorithms that implement ‘one-shot learning’ [1, 2], the neural underpinnings of this abrupt form of learning have remained elusive. In the hippocampus, a region recognized for its importance in learning and memory, behavioral timescale synaptic plasticity (BTSP) has emerged as a robust mechanism for the rapid generation of spatial representations (place fields) following the occurrence of plateau potentials [3–12]. However, the hippocampus not only represents the location of animals in space [13–16], but also non-spatial sensory information [17–20]. The hippocampus dynamically links these sensory experiences across time through sequential firing that tracks the passage of time after specific events [15, 20–23]. It is unclear whether BTSP also plays a role in the formation of non-spatial sensory-driven or internally generated hippocampal representations, which can form the basis for ‘one-shot learning’.

Hebbian plasticity mechanisms such as spike-timing dependent plasticity require causality and many repetitions to potentiate synapses when presynaptic spikes precede postsynaptic action potentials by a few milliseconds [24–27]. While this mechanism may play a role in modulating hippocampal responses, BTSP has many features which could make it a more robust and rapid mechanism for the generation of non-spatial receptive fields. During spatial learning tasks, a single calcium plateau potential can serve as the induction event, asymmetrically boosting synaptic inputs that occur up to several seconds before the induction event, leading to a membrane potential ramp and reliable spatial firing on subsequent trials [4, 7, 8]. It is not known whether plateau potentials occurring during non-spatial tasks could also boost synaptic inputs at specific timepoints in the task leading to the rapid formation of stable representations of sensory stimuli and time. The rapid induction of these non-spatial hippocampal representations by BTSP could form the basis for rapid learning in the hippocampus.

The role of the entorhinal cortex (EC) in inducing BTSP events [9] and relaying sensory information during non-spatial tasks [19, 28, 29] is poorly understood. CA1 receives direct layer III EC input via the temporammonic (TA) pathway and indirect input via the perforant path from layer II EC to dentate gyrus (DG). DG then projects to CA3, which in turn projects to CA1 [30, 31]. Lateral and medial EC (LEC and MEC) have distinct inputs and behaviorally relevant response properties: LEC robustly represents olfactory information [32–35], while MEC mainly encodes visuo-spatial information [36–39]. Furthermore, the MEC plays a major role in the induction of plateau potential ‘teaching signals’ during BTSP induced during spatial learning tasks [9]. Yet, whether MEC and LEC play distinct roles in the generation of BTSP during non-spatial tasks remains to be determined.

To address these questions, we investigated multimodal representations within CA1 and EC during a non-spatial olfactory delayed non-match-to-sample (DNMS) working memory task [20]. We have previously shown that CA1 pyramidal neurons fire sequentially in response to specific odors and across the 5-second delay period during DNMS performance [20]. We hypothesized that non-spatial BTSP can generate odor representations in CA1 and that this process would be modulated by MEC and LEC inputs. Using two-photon calcium imaging of dorsal CA1, we recorded non-spatial ‘BTSP-like’ events that formed odor-specific fields in CA1 during expert performance of the DNMS task. Combining this approach with holographic optogenetic stimulation, we were able to successfully generate odor-specific representations in individual cells following a single stimulation. Recording dorsal CA1 across learning of the task revealed higher rates of BTSP-like events during the first day of learning compared to all subsequent days. In addition, we found that MEC chemogenetic inhibition decreased the frequency of large calcium induction events in CA1, while LEC inhibition reduced the success rate of odor-field generation. Finally, two-photon calcium imaging of temporammonic EC axonal projections to CA1 demonstrated that MEC axonal strong firing to odor presentations likely drives large plateau-like calcium events in CA1, while LEC axonal odor-selectivity mediates plasticity in the formation of odor-fields after the large calcium induction event. Therefore, non-spatial BTSP leads to the generation of odor-fields during working memory performance and is strongly modulated by entorhinal cortex.

## RESULTS

We used *in vivo* two-photon calcium imaging to record the activity of neurons in the pyramidal layer of dorsal CA1 while animals performed an olfactory delayed non-match-to-sample (DNMS) working memory task (**Fig. 1a-e**). Adult male and female mice (n=17) were injected with AAV1-Syn-jGCaMP8f into the right dorsal CA1 and implanted with a 3mm diameter glass-bottomed titanium cannula above the intact alveus after aspiration of the overlying cortex and corpus callosum [20] (**Fig. 1d-e**). After one week of recovery, mice were water-restricted and trained on the olfactory DNMS working memory task [20, 40], while head-fixed on a spherical treadmill (**Fig. 1a-b**). Each trial consisted of two 1-second odor presentations separated by a 5-second delay period. One second after the offset of the 2^nd^ odor, there was a 3-second reward period during which the choice of the animal was determined. Mice were trained to lick the lickport to release water during this reward period if the two odors did not match (correct ‘hit’). Mice learned to refrain from licking the lickport if the odors matched (correct ‘rejection’), and overall performance was quantified as the percentage of correct ‘hits’ and correct ‘rejections’ out of all trials. We considered performance above 85% to be expert level. Each session of the DNMS task consisted of 5 blocks of 20 trials, with pseudorandomly distributed odor combinations (**Fig 1c**). Mice were recorded for 8 days during expert performance for a total of 136 recording sessions yielding an average of 312 ± 125 (mean ± standard deviation) active neurons per day. We successfully imaged the same field of view (FOV) for each of the 8 days for all animals. Calcium signals were extracted and deconvolved using Suite2p [41] (see methods).

**Figure 1:**
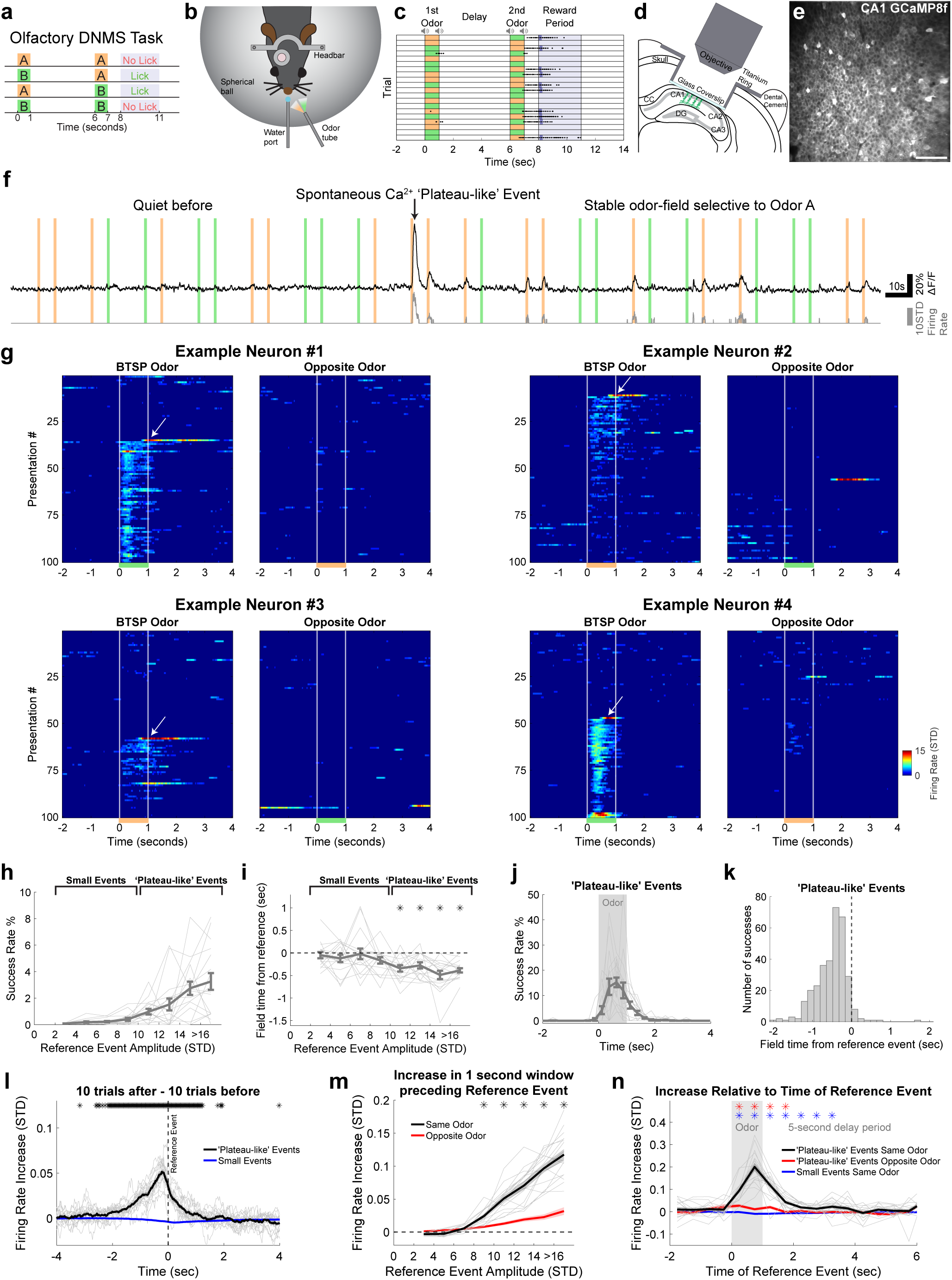
Behavioral timescale synaptic plasticity (BTSP) in a non-spatial working memory task. **a,** Schematic of the olfactory delayed non-match-to-sample (DNMS) task. Water delivery and licking behavior was assessed during the 3-second reward period. **b,** Mice were head-fixed above a Styrofoam spherical ball to allow running. **c,** Example session of perfect performance for 20 trials. Dots indicate licks and dark blue bars indicate water delivery. **d,** Schematic of two-photon calcium imaging of dorsal CA1 pyramidal neurons. **e,** Example field of view of CA1 neurons expressing GCaMP8f. Scale bar is 100µm. **f,** Example trace of one neuron with a ‘BTSP-like’ event and odor-field formed. Colored bars indicate odor presentations. Black trace is ΔF/F, and gray is z-scored deconvolved signal. **g,** Four examples of ‘BTSP-like’ events. White vertical lines indicate odor onset and offset, and white arrows point to spontaneous induction ‘plateau-like’ events. **h,** Success rate of a calcium event generating an odor-field increases with induction-event amplitude. Success rate is defined as the percentage of events that generate a significant odor-field. Individual thin gray lines represent each of the 17 animals. Error bars represent the standard error of the mean across the animals. Events above 10 STD are considered ‘plateau-like’ events. **i,** Asymmetrical field formation with trial-time difference between ‘plateau-like’ event peak and formed odor-field peak. This difference is only significant for ‘plateau-like’ events. Statistics are one-sample t-tests, and asterisks indicate bins with p < 0.05 (corrected for multiple comparisons using the Benjamini-Hochberg procedure). **j,** Success rate is highest during odor presentation (for only ‘plateau-like’ events). **k,** Histogram showing asymmetrical distribution of formed-fields for all 323 successful ‘plateau-like’ events. **l,** Firing rate increase calculated as the average of 10 same odor trials after reference event subtracted by 10 same odor trials before reference event. Individual thin gray lines represent the average of all ‘plateau-like’ events aligned to the reference event for each of the 17 animals. Thick black line represents the mean, and shading represents the standard error. Blue line and shading represent the same calculation for all small events with amplitude less than 5 STD. Asterisks indicate bins with p < 0.05 **m,** Calculated as the average of the ramp from (l) between −1 and 0 seconds in trial-time, plotted as a function of reference event amplitude. Gray lines and black line show individual animals and mean for the increase in same odor trials. Red is the same calculation for 10 trials of the opposite odor after and 10 trials of the opposite odor before. Statistics are paired t-tests comparing same and opposite odor measurements, and asterisks indicate bins with p < 0.05 (corrected for multiple comparisons using the Benjamini-Hochberg procedure). **n,** Same calculation as in (m), but as a function of reference event timing in trial-time (excluding time after delay period). Blue asterisks represent significance in paired t-tests comparing the black and blue traces, while red asterisks represent significance in comparing black and red traces.

### Non-spatial BTSP-like events in CA1 formed stable odor-specific fields

In our previous work, we found that a population of CA1 hippocampal neurons fired in an odor-specific manner during specific epochs of the DNMS task; some fired during the presentation of specific odors (‘odor fields’) and others at timepoints during the delay period (‘time fields’) after presentation of specific odors [20]. Here, we observed CA1 neurons with activity patterns consistent with BTSP during expert DNMS performance (**Fig. 1f-g and Supp. Fig. 1 and 2**). Namely, neurons without a clear odor or time field developed a stable field after a single spontaneous large calcium event.

A single calcium ‘reference event’ was considered an induction event that generated an odor-field using a set of strict criteria (see methods). With increasing ‘reference event’ amplitude, the success rate for induction of an odor-field also increased (**Fig. 1h**), suggesting that these induction events drive the formation of odor-fields. Only events larger than 10 standard deviations (STD) of the mean deconvolved signal yielded a significantly asymmetrical field-formation with the new odor-field occurring earlier in trial-time on subsequent trials compared to their reference event (**Fig. 1i**); this pattern of field induction was very similar to spatial BTSP recorded in other studies [3–12]. We therefore considered a reference calcium event as ‘plateau-like’ when its amplitude was greater than 10 STD. For ‘plateau-like’ events that were preceded by minimal activity and no significant field, the probability of field-induction peaked during odor presentation or immediately after odor offset, with the average success rate of field formation peaking at 15% during the second half of the odor presentation period (**Fig. 1j**). Only 26 ‘plateau-like’ events (8% of the 323 successful events) yielded time-fields beyond 0.5 seconds after odor offset. The newly formed odor-specific fields peaked 0.42 ± 0.14 seconds earlier in trial-time compared to the reference event timepoint (n=323 successful ‘plateau-like’ events) (**Fig. 1k**). The small events that represented the random chance of an event passing our strict criteria had a success rate that only peaked at 0.6% during odor presentation, and they did not yield significant asymmetrical field-formation (**Supp. Fig. 3**).

Categorizing BTSP-like events into successes and failures restricted analysis to a narrowly defined type of plasticity event based on formation of significant odor responses. However, some responses were reinforced by multiple ‘plateau-like’ events (**Supp. Fig. 2**). Therefore, we developed a simpler analysis to measure potentiation as the increase in firing rate across 10 trials after versus 10 trials before a reference calcium event (‘firing rate increase’). We observed an asymmetrical ramp of firing rate increase with a peak in subsequent trials 0.25 seconds earlier in trial-time than the reference event timepoint (**Fig. 1l**). No increase was observed when aligned to small events (**Fig. 1l**). While the firing rate increase was significant up to 3 seconds earlier and 1 second later in trial-time, we focused on the average increase in the 1-second period preceding the reference event. Similar to the previous binary success rate analysis, we found that a significant firing rate increase only occurred after events greater than 8 STD (**Fig. 1m**). This increase was highly odor selective (**Fig. 1m**). Firing rate increase peaked during the second half of the odor presentation and remained significant up to halfway through the delay (**Fig. 1n**), suggesting that time-fields during the delay period may form after several successive plateau potentials. Results were similar for a range of trial numbers averaged before and after the reference event (**Supp. Fig. 4a-d**). Both analysis methods demonstrate non-spatial BTSP leading to the formation odor-fields characterized by an asymmetric firing rate increase ramp.

To determine if locomotion influenced non-spatial BTSP-like events, we recorded treadmill locomotion during the task. Mice exhibited a range of locomotion patterns while performing the task with some mice rarely moving on the treadmill, some primarily flinching or twitching at the onset of some odor presentations, and others with bouts of running throughout the trial. On average, minor jitter movements occurred at the onset of 20% of odors and running occurred in up to 5% of trials (**Supp. Fig. 5a-b**). On average, animals spent 90.4 ± 2.3 % of the time immobile on the treadmill, 7.0 ± 1.7 % of the time with minor jitter movements, and only 2.6 ± 1.8 % of the time running (**Supp. Fig. 5c**). As expected [42–44], movement increased neural activity. Both minor jitter movements and running increased the rate of small events and ‘plateau-like’ events (**Supp. Fig. 5d-e**). However, neither of these types of movements during reference ‘plateau-like’ events influenced the success rate of field formation or firing rate increase (**Supp. Fig. 5f-g**). In total, out of the 323 detected successful events, 235 (73%) occurred during periods without movement, 70 (22%) occurred during minor jitter movements, and only 18 (6%) occurred during running bouts. Moreover, mice spent 13.9 ± 4.2 % of time licking (primarily during the reward period of non-match trials) (**Supp. Fig 5h-j**), which also increased the rate of ‘plateau-like’ events (**Supp. Fig. 5k-l**). Yet, the success rate and firing rate increase was lower for licking bouts (**Supp. Fig. 5m-n**). In total, 289 (89%) occurred during periods without licking, and only 34 (11%) occurred during licking bouts. Collectively, our findings reveal non-spatial BTSP forming odor-specific fields that are not driven by locomotion or licking behavior.

### Single-neuron holographical optogenetic stimulation induced odor-fields

To confirm that large calcium events causally drive the formation of odor-specific fields, we performed simultaneous two-photon calcium imaging and single-neuron holographic optogenetic stimulation in three animals genetically expressing GCaMP (two mice GCaMP6s and one mouse GCaMP8s). Mice were injected with AAV8-CaMKIIa-ChRmine-mScarlet-Kv2.1-WPRE into the right dorsal CA1 to drive expression of the excitatory opsin ChrRmine in pyramidal neurons (**Fig. 2a**). Individual CA1 neurons of well-trained mice were stimulated for one second (10ms pulses at 10Hz) at various timepoints throughout the entire trial period from 1^st^ odor presentation to middle of the reward period. After recording at least 4 baseline trials, individual stimulation points were delivered at least 6 seconds apart during task performance. Across multiple days and different FOVs with up to 72 stimulated neurons per day, we successfully induced ‘BTSP-like’ events yielding new stable odor-fields following a single stimulation (**Fig. 2b-c**). Induced event amplitudes varied in size (likely due to varying levels of ChRmine expression and/or excitability state of the neuron) (**Fig. 2d**), but all successful induction events were greater than 10 STD, validating our previously used criteria. Importantly, the success rate of field formation and firing rate increase after stimulation (**Fig. 2e-f**) were similar to the spontaneous events we recorded in Figure 1. Therefore, large plateau-like events causally drive the formation of odor-fields.

**Figure 2:**
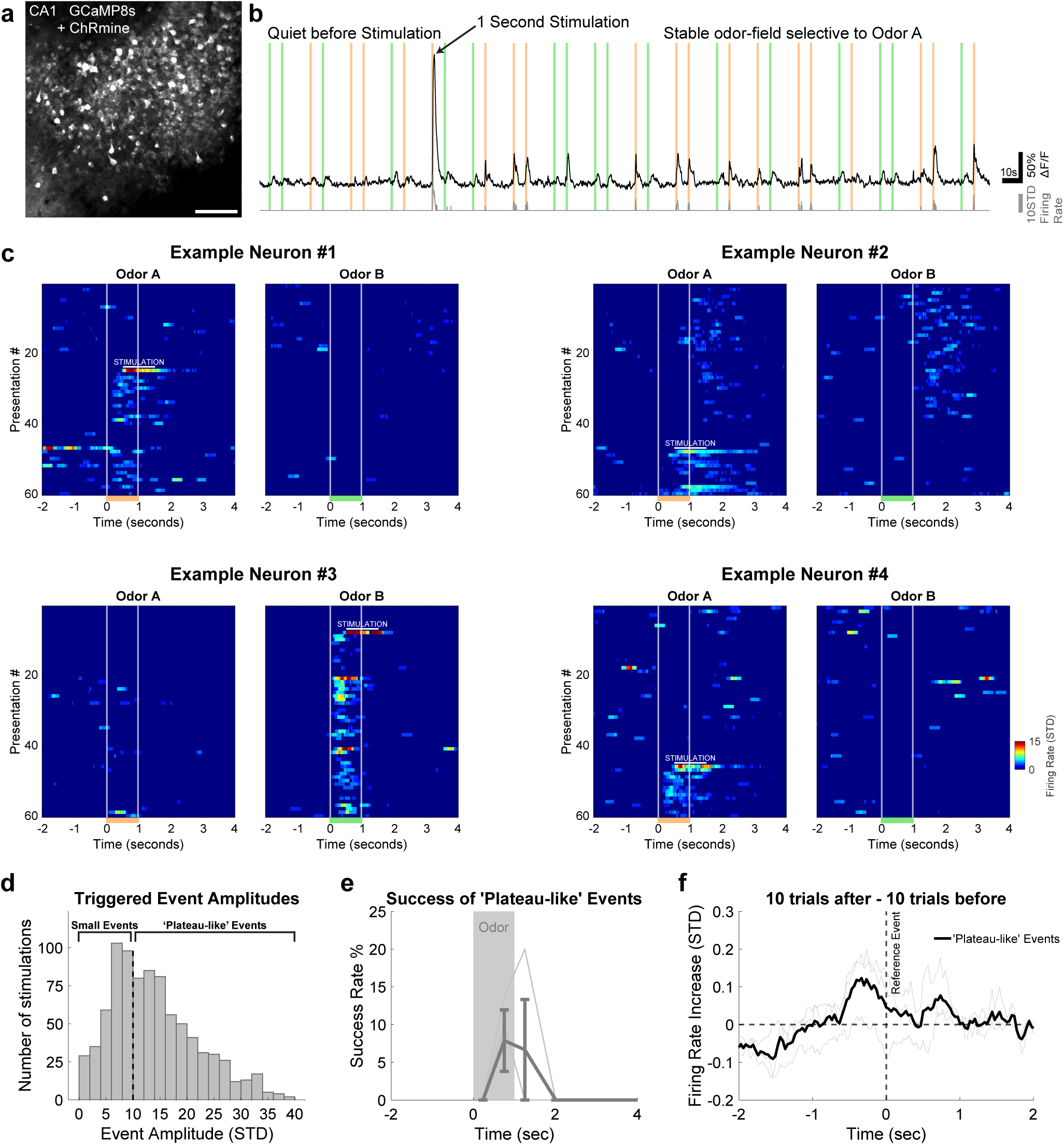
Single-neuron holographical optogenetic stimulation induced odor-fields. **a,** Example field of view of CA1 neurons expressing GCaMP8s and ChRmine. Scale bar is 100µm. **b,** Same as Fig. 1f, but for a single neuron responding to optogenetic stimulation with a large calcium event followed by a stable odor-field. **c,** Same as Fig. 1g, but for 4 neurons that received one-second optogenetic stimulation. White bar above large calcium event indicates the timing of stimulation. **d,** Histogram shows distribution of all triggered event amplitudes (n=857). **e,** Same as Fig. 1j, but for the three animals with optogenetic stimulation and only including stimulated events above 10 STD. **f,** Same as Fig. 1l, but for the three animals and stimulated events above 10 STD. Note that noisier appearance of data compared to Fig. 1l is likely due to much fewer events to average per animal.

### Highest rates of non-spatial BTSP occurred early in learning

Given the role of spatial BTSP in formation of place cells during learning of a novel spatial environment, we next asked if non-spatial BTSP-like events were also more frequent earlier in learning of non-spatial cues. To address this question, we repeated two-photon calcium imaging of pyramidal CA1 neurons in five mice during learning of the olfactory DNMS task. To include learning of the odor cues themselves, we introduced them only at the final training stage. Mice were first trained to lick at the appropriate reward window by providing only auditory clicks at the timepoints used for odor onset and offset. This allowed the mice to learn the temporal structure of the task without receiving any odors (**Fig. 3a**). Mice licked during the reward period for more than 90% of trials during these ‘odor-free’ shaping days. On the following ‘Day 1’, odors were delivered for the first time and mice continued to lick during most trials yielding chance-level performance on Days 1 and 2. By days 3 and 4, mice began refraining from licking on match trials (**Fig. 3b**) and gradually increased their performance to expert-level (85%) around Day 5 (**Fig. 3c**). Imaging was conducted on the last two days of ‘odor-free’ shaping (Days −2 and −1), and 9 days during the final stage of the olfactory DNMS task. The same FOV was used on each day for each animal.

**Figure 3:**
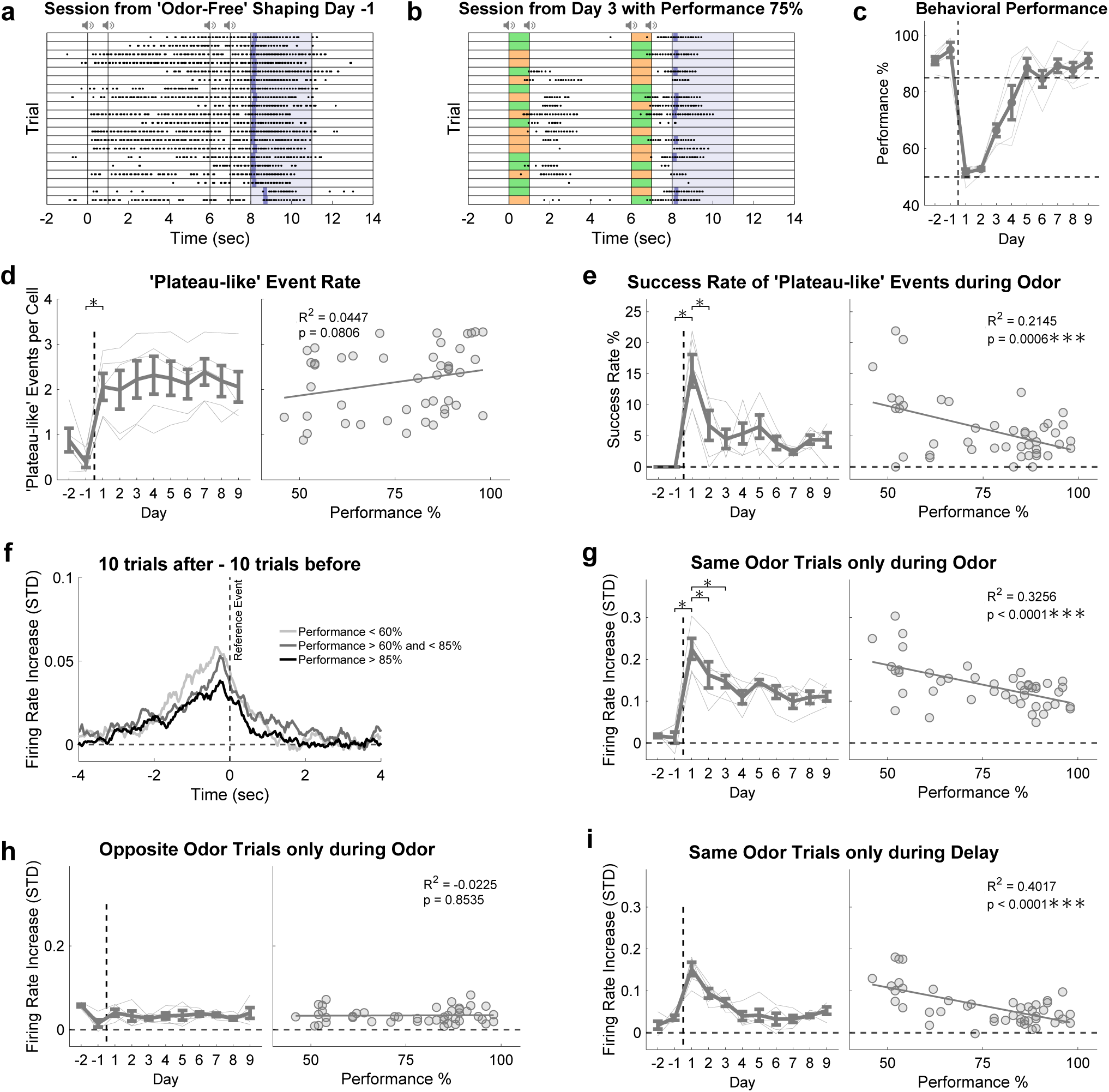
Highest rates of non-spatial BTSP occurred early in learning. **a,** Example session of 20 trials during ‘odor-free’ shaping Day −1. The gray speaker icons indicate time of auditory clicking. Dots indicate licks and dark blue bars indicate water delivery. **b,** Same as Fig. 1c, but for a session from Day 3 with 75% performance. **c,** Behavioral performance across all 11 days of imaging. Vertical dashed lines represent the transition from baseline ‘odor-free’ shaping to the standard olfactory DNMS task. Horizontal dashed lines represent chance-level performance at 50% and expert-level performance at 85%. Individual thin gray lines represent each of the 5 animals. Error bars represent the standard error of the mean across the animals. **d,** Number of events greater than 10 STD per neuron per day. Left panel asterisk shows significance of difference between Day −1 and Day 1 (paired t-test, p = 0.016). Right panel shows correlation with performance. Statistics are from multiple linear regression with mixed-effects (fixed effect of performance and random effect of animal). **e,** Same as (d) but for success rate of forming an odor-field only when the ‘plateau-like’ event occurred during the odor presentation. Asterisks in left panel (paired t-tests, p = 0.046 and p = 0.018) **f,** Same as Fig. 1l showing firing rate increase ramp for 3 different levels of performance. Each line is the average of the 5 animals. **g,** Average firing rate increase during 1 second window preceding reference event in trial-time for ‘plateau-like’ events that occurred during odor presentation. Asterisks in left panel (paired t-tests, p = 0.005, p = 0.008, p = 0.047) **h,** Same as (g), but for 10 opposite odor trials after and before. **i,** Same as (g), but for ‘plateau-like’ events that occurred during the delay period.

Compared to baseline ‘odor-free’ shaping days, the ‘plateau-like’ event rate strongly increased in Day 1 and remained stable across all subsequent days of learning, without correlation with performance (**Fig. 3d**). However, the success rate of these ‘plateau-like’ events generating an odor-field peaked on Day 1 compared to the following days and was strongly anti-correlated with performance (**Fig. 3e**). Analysis of the firing rate increase ramps (**Fig. 3f**) revealed a similar effect when compared across the days of learning for the same odor trials (**Fig. 3g**). Notably, the very small firing rate increase during the opposite odor trials remained stable through all days of baseline and learning (**Fig. 3h**), suggesting that these non-specific increases may be due to other factors such as inputs related to auditory clicks rather than odor-specific information. Smaller firing rate increases observed during the delay period also showed a strong negative correlation with performance for same odor trials (**Fig. 3i**). Importantly, minor jitter movements and running did not change significantly across learning (**Supp. Fig. 6**), thus they cannot account for differences observed across days. Therefore, despite a stable rate of ‘plateau-like’ events during learning, the rate of non-spatial BTSP-like events was highest early in learning of novel odor cues.

### Chemogenetic inhibition of entorhinal cortex disrupted non-spatial BTSP

Entorhinal inputs can drive BTSP induction events during spatial learning tasks [9, 45]. To determine whether entorhinal inputs may also play a role in the generation of ‘plateau-like’ events during non-spatial BTSP, we used chemogenetic inhibition of lateral entorhinal cortex (LEC) or medial entorhinal cortex (MEC), while imaging CA1 calcium activity during expert performance of the working memory task. Mice were injected with AAV1-Syn-jGCaMP8f in the dorsal CA1 and were subsequently implanted with an optical canula over CA1 as in the previous section. These mice also received an injection of AAV5-CaMKII-PSAM4 into either LEC (n=6 mice) or MEC (n=5 mice) to express the potent chemogenetic inhibitor PSAM4 [46] in excitatory neurons of either structure (**Fig. 4a**). Control mice underwent injections of AAV5-CaMKII-mCherry into either LEC (n=3 mice) or MEC (n=3 mice). Animals were water-restricted, trained on the task, and imaged three weeks after viral expression. Each animal was recorded for eight days after reaching expert level performance. Between 10-20 minutes before two-photon calcium imaging began on each day, mice received an intraperitoneal (IP) injection of saline or uPSEM (the effector molecule for PSAM4). Saline and uPSEM injections were alternated daily and animals were counter-balanced such that one half of the mice received injections of uPSEM on the first day and the other half received injections of saline. We compared the activity of matched neurons over four pairs of saline and uPSEM days during expert performance when BTSP-like event rates were stable.

**Figure 4:**
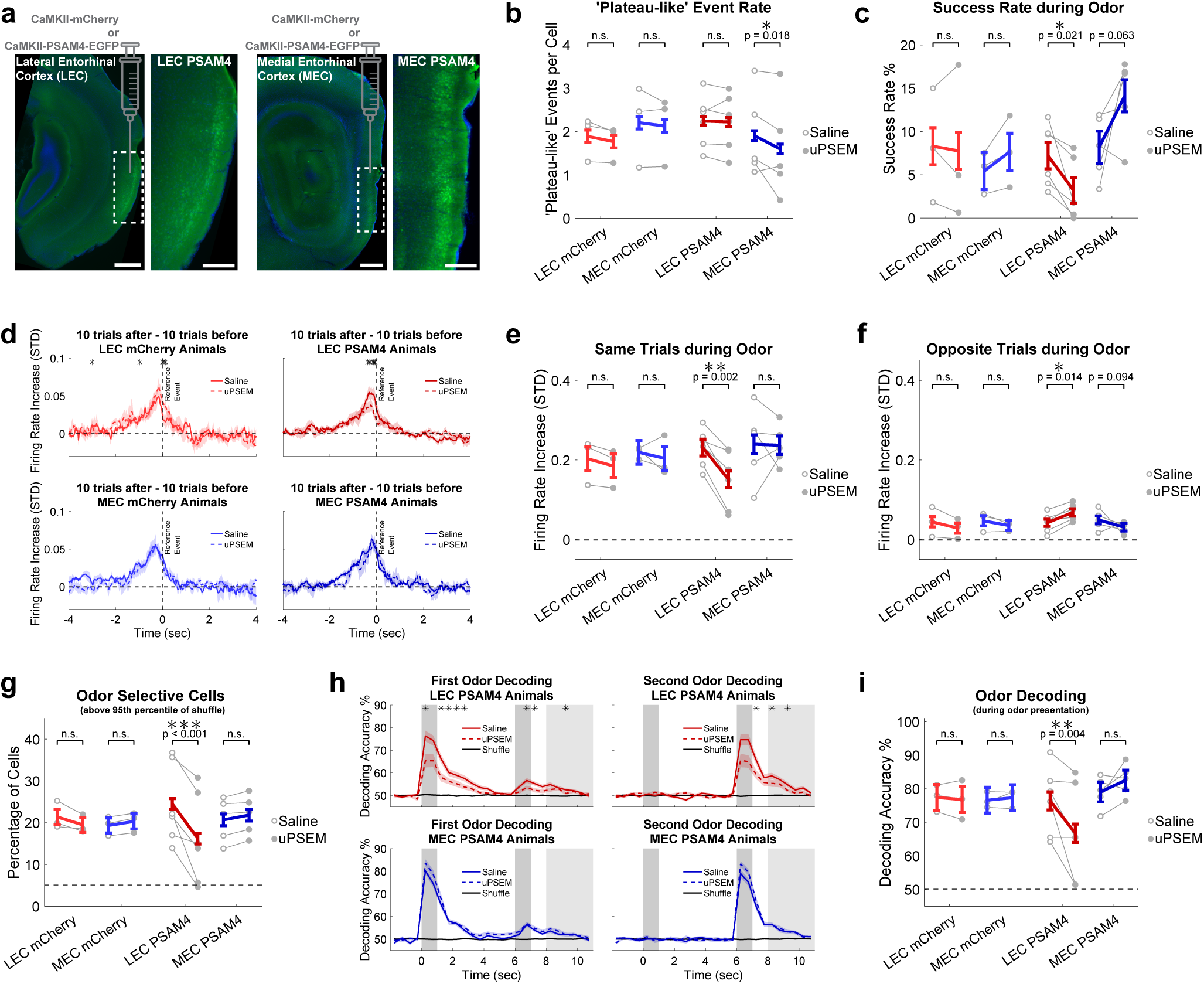
LEC and MEC inhibition differentially modulated BTSP, and LEC inhibition weakened odor-selectivity in CA1. **a,** Injections of virus to drive the expression of mCherry or PSAM4 were delivered to either LEC or MEC. Images showing LEC are from coronal sections, while MEC are from sagittal sections. For both LEC and MEC, the larger image on the left has a 500µm scale bar and the right image is a zoom of the white outline with a 200µm scale bar. **b,** Number of events greater than 10 STD per neuron per day. Paired dots show animal averages representing the pairs of imaging days (averaging 4 pairs per animal). Statistic is a three-way ANOVA (group, pair, and day-type; within-subject for pair and day-type, between-subject for group; multiple comparisons corrected with Tukey-Kramer test). **c,** Success rate of ‘plateau-like’ events generating an odor-field. Statistics are the same as (B). **d,** Same as Fig. 1l showing firing rate increase ramp difference between saline and uPSEM in each animal group. Lines represent animal averages, and shaded areas represent the standard error. Statistics are a three-way ANOVA (pair, day-type, and bin; within-subject for all 3; multiple comparisons corrected with Tukey-Kramer test). Asterisks indicate bins with p < 0.05. **e,** Same calculation as Fig. 3g but plotted the same as panels above. **f,** Same calculation as Fig. 3h. **g,** Percentage of neurons that had a selectivity value above 95^th^ percentile of shuffle. **h,** Binary support vector machine (SVM) decoding of first and second odor across the trial structure with 0.5 second bins for experimental animal groups (repetitive subsampling of 100 neurons for each recording session). Thinner gray bars indicate odor presentation and wider bar from seconds 8-11 is the reward period. Statistics are same as (d). **i,** Odor decoding performance only during the odor presentation period.

Despite a lack of a behavioral effect (**Supp. Fig. 7a**), both LEC and MEC inhibition strongly affected the non-spatial ‘BTSP-like’ events. MEC, but not LEC, inhibition significantly reduced the number of ‘plateau-like’ events from 1.91 ± 0.98 per neuron per day to 1.60 ± 1.12 per neuron per day (**Fig. 4b**). In contrast, LEC, but not MEC, dramatically reduced the success rate of ‘plateau-like’ events during odor presentations from 7.20 ± 3.61% to 3.18 ± 3.48% (**Fig. 4c**). Additionally, only LEC inhibition significantly decreased the firing rate increase ramps during the 1-second window preceding reference ‘plateau-like’ events on uPSEM days (**Fig. 4d**). For ‘plateau-like’ events during odor presentation of LEC PSAM animals, the firing rate increase was significantly lower on uPSEM days for same odor trials but significantly higher for opposite odor trials (**Fig. 4e-f**). Importantly, neither LEC nor MEC inhibition affected minor jitter movements, running, or licking, so these effects could not be explained by differences in animal movement (**Supp. Fig. 7b-e**). Together, these findings suggest that MEC affects the generation of large ‘plateau-like’ events in CA1, while LEC activity increases the likelihood that these events result in successful field generation. The differential effects of firing rate increase between same odor and opposite odor trials strongly suggest that LEC inputs to the hippocampus are necessary for relaying odor-specific information.

### LEC inhibition reduced strength of odor representations in dorsal CA1

Given that LEC inputs have been previously shown to encode odor-related information [32–35], we hypothesized that they could convey odor-related information to CA1 in our DNMS task. If so, we would expect the inhibition of LEC but not MEC to decrease odor-selectivity in CA1, potentially driving the decrease in success rate of ‘plateau-like’ events in generating odor-fields. Indeed, LEC chemogenetic inhibition significantly decreased the percentage of significantly odor-selective neurons from 24.5 ± 9.6% on saline control days and only 16.2 ± 10.8% on uPSEM inhibition days (**Fig. 4g**), while having no effect on any of the other animal groups. To further confirm this effect, we performed binary support vector machine (SVM) decoding training and testing on the same day to evaluate the relative strength of odor encoding on saline days compared to uPSEM days subsampling only 100 neurons. Overall, LEC inhibition significantly decreased odor decoding accuracy, while MEC inhibition did not alter it (**Fig. 4h**). While the effect in LEC experimental PSAM4 animals also was observed during the first half of the delay period, the effect during the odor presentation period was the strongest with decoding accuracy at 76.4 ± 10.2% on saline control days and only 66.7 ± 14.3% on uPSEM inhibition days. Odor decoding of MEC experimental PSAM4 and control animals expressing mCherry was unaffected by uPSEM administration. Collectively, LEC inhibition strongly decreased odor-selectivity and decodability in pyramidal CA1 neurons, which may have driven the reduction in the success rate of ‘plateau-like’ events in generating odor-fields, though additional mechanisms could potentially contribute to this effect.

### Two-photon calcium imaging of temporammonic entorhinal cortical axons in dorsal CA1 revealed differential sequential activity in LEC and MEC inputs

The EC is the primary cortical input to the hippocampus; CA1 receives direct layer III EC input via the temporammonic (TA) pathway and indirect input via the perforant path from layer II EC to dentate gyrus, which then projects to CA3, which in turn projects to CA1 [30, 31]. Given the contribution of MEC in driving ‘plateau-like’ events, we asked if there are differences in timing of LEC and MEC TA inputs. Also, given the strong differences in odor decodability observed in dorsal CA1 with LEC versus MEC inhibition, we asked whether TA inputs from LEC and MEC differ in the olfactory information they convey to CA1.

To address these questions, we performed two-photon calcium imaging of LEC or MEC TA axons in dorsal CA1 during expert performance of the DNMS task. Adult male and female mice were injected with AAV1-CaMKII-Cre and AAV1-CAG-FLEX-jGCaMP7s in either LEC (n=8) or MEC (n=8) (**Fig. 5a,c**). Mice were implanted with hippocampal windows as in the previous experiments. After three weeks of expression, there was extensive GCaMP7s axonal expression of TA inputs within the stratum lacunosum-moleculare (SLM) layer, as well as layer II EC perforant path axons ramifying deeper within the stratum moleculare (MOL) layer of the dentate gyrus (**Fig. 5a,c**). *In vivo*, we were able to selectively image TA EC axons 300-400 µm beneath the alveus. Post-hoc histology after two-photon imaging experiments also confirmed that all mice had extensive expression of GCaMP7s in axons within the SLM layer of hippocampus and somatic expression restricted to either LEC or MEC. Recordings were processed with Suite2p [41] using parameters optimized for axonal imaging, followed by post-hoc fusion of axon segments with highly correlated activity which were branches of the same axon (see methods) (**Fig. 5b,d**). Only behavioral sessions with expert-performance, with no difference between the groups, were used for the following analysis (**Supp. Fig. 8c**).

**Figure 5:**
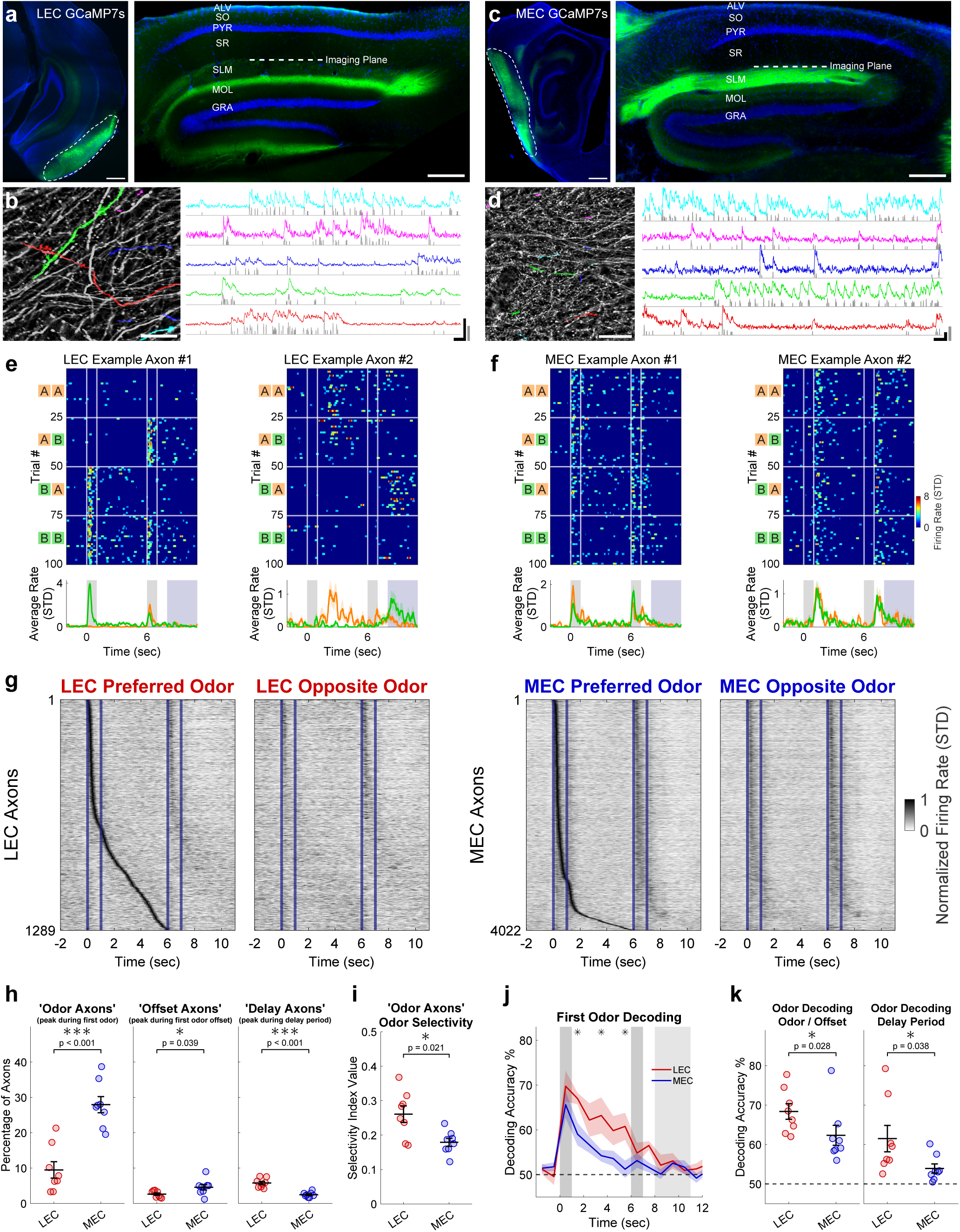
Two-photon calcium imaging of temporammonic entorhinal cortical axons in dorsal CA1 revealed differential sequential activity in LEC and MEC inputs and stronger odor-selectivity of LEC axons. **a,** Coronal sections showing GCaMP7s expression LEC (left panel, scale bar = 500µm) and in dorsal hippocampus (right panel, scale bar = 250µm). Blue is DAPI. Imaging plane is at the superficial part of the SLM layer, which is the first layer of axons visible when lowering into the tissue roughly 300-400µm beneath the coverglass. ALV = alveus, SO = stratum oriens, PYR = stratum pyramidale, SR = stratum radiatum, SLM = stratum lacunosum-moleculare, MOL = stratum moleculare, GRA = stratum granulare. **b,** Field of view from the same animal (scale bar = 50µm), with 5 example masks and their corresponding fluorescence traces. Gray is z-scored deconvolved signal. Black horizontal scale bar = 10 seconds. Black vertical scale bar = 5% ΔF/F. Gray vertical scale bar = 10 STD normalized deconvolved signal. **c,d,** Same as (a,b) but for MEC and showing sagittal sections. All scale bars are the same. **e,** Two example axon segments showing odor-specific firing. The left axon had its peak during the odor presentation, while the right one had its peak during the delay period. Heatmaps show deconvolved signals on each trial that was grouped into trial type. Average traces at bottom show difference in average firing rate split by trials that started with Odor A and those that started with Odor B. **f,** Same as (e) but for two representative MEC axon segments with less odor-selectivity. The right axon had its peak following the offset of the odor presentation. **g,** Sequential activity of only axon segments that had a significant peak during the first odor presentation or delay period. Each row is the average trace of trials (normalized to peak on preferred odor trials and sorted by order of peak in trial-time). Opposite odor panels show the axons in the same rows on trials of their non-preferred odor trial type. Blue lines indicate odor onset and offset. **h,** Percentage of axons with a significant peak during the first odor presentation (‘odor axon’ trial-time 0-1), odor offset (‘offset axons’ 1-2 trial-time), and delay period (‘delay axons’ 2-6 trial-time). Each circle represents the animal average of expert-level performance recording sessions. Statistics are independent samples t-test of animal averages. **i,** Average odor-selectivity values of all ‘odor axons’. Statistics are the same as (h). **j,** Binary support vector machine (SVM) decoding of first odor using 10 most odor-selective neurons from each recording session (bin size of 1 second). Statistics are independent samples t-tests, and asterisks indicate bins with p < 0.05 (corrected for multiple comparisons using the Benjamini-Hochberg procedure). **k,** Average decoding during the 2 bins from timepoints 0-2 seconds, and then the 4 bins from timepoints 2-6 seconds. Left panel statistics are independent t-test p = 0.081; two-sides Wilcoxon rank sum test p = 0.028 displayed because of outlier. Right panel statistics are independent t-test p = 0.050; two-sided Wilcoxon rank sum test p = 0.038.

A proportion of LEC and MEC TA axons responded reliably at specific timepoints with some having significant peaks during the odor presentation, some during odor offset, and others during the delay period (**Fig. 5e-f**). Altogether, the LEC or MEC axonal populations had sequential activity that tiled the entire first odor presentation and delay period. However, these sequences differed drastically between LEC and MEC (**Fig. 5g and Supp. Fig. 8a-b**). A much higher percentage of MEC axons had significant peaks during odor presentations and their offset compared to LEC axons (MEC with 28.0 ± 6.5% ‘odor axons’ compared to only 9.5 ± 6.5% for LEC; MEC with 4.6 ± 2.2% ‘offset axons’ compared to only 2.7 ± 1.0% for LEC) (**Fig. 5h**). MEC ‘odor axons’ also showed greater trial reliability compared to LEC (**Supp. Fig. 8d**). In contrast, a higher percentage of LEC axons had significant peaks during the delay period (LEC with 5.8 ± 1.3% ‘delay axons’ compared to only 2.4 ± 0.7% for MEC). Given that MEC inhibition reduced the rate of ‘plateau-like’ events in CA1, we hypothesize that this strong MEC input timed to the odor presentation is likely important for driving ‘plateau-like’ events.

### LEC temporammonic axons carried strong odor-specific information than MEC axons

Although a higher proportion of MEC axons were tuned to firing during the odor presentation, examination of sequential firing patterns suggested that these axons were firing with less odor-specificity (**Fig. 5g and Supp. Fig. 8a-b**). LEC axons with a significant peak during the odor presentation (‘odor axons’) had larger odor-selectivity values than MEC (LEC with an average of 0.261 ± 0.068 compared to only 0.179 ± 0.033 for MEC) (**Fig. 5h**). Again, to confirm this difference in odor-selectivity, we performed binary support vector machine (SVM) decoding training and testing on the same day using only the most selective 10 axons from each recording session. First odor decoding accuracy was higher in LEC compared to MEC for both the odor/offset period and the remaining 4 seconds of the delay period (68.4 ± 5.6% for LEC and 62.3 ± 7.2% for MEC during odor/offset 0-2 second window; 61.5 ± 9.5% for LEC and 53.9 ± 3.3% for MEC during odor/offset 2-6 second window). Altogether, these findings suggest that LEC temporammonic axonal odor representations were stronger than MEC axonal odor representations, and this may explain the decrease in success rate and firing rate increases with LEC inhibition, although strong odor information is also likely being relayed indirectly through the CA1 perforant path as well.

## DISCUSSION

Using two-photon calcium imaging of dorsal CA1 pyramidal neurons during an olfactory working memory task, we find that non-spatial representations can form on single trials following large calcium events. These events have characteristics of behavioral timescale synaptic plasticity (BTSP), such as asymmetrical formation, reported previously during spatial tasks [3–12], suggesting that BTSP may be a general plasticity mechanism for formation of hippocampal representations during both spatial and non-spatial cognition. Holographic optogenetic stimulation confirms causality of single large calcium events driving the rapid formation of odor representations. Spontaneous BTSP-like events are much more frequent on the first day of task learning and anti-correlated with performance, suggesting that BTSP may be important for early learning of non-spatial cues. Additionally, MEC and LEC inhibition differentially modulate non-spatial BTSP during working memory performance. MEC inhibition decreases the frequency of large ‘plateau-like’ calcium events, while LEC inhibition reduces potentiation and the success rate of these ‘plateau-like’ events generating an odor-field. LEC inputs are critical for the formation of odor representations in CA1, with LEC inhibition dramatically weakening CA1 odor selectivity and odor encoding. This may contribute to their modulation of BTSP success rate in generating odor-selective responses. Finally, performing two-photon calcium imaging of LEC or MEC temporammonic pathway axons to CA1, we find that MEC’s strong firing to odor presentations likely drives large plateau-like calcium events in CA1, while LEC’s stronger odor-selectivity is likely necessary for relaying of odor information to the hippocampus to allow odor-specific potentiation of synapses.

This is to our knowledge the first description of BTSP occurring in a non-spatial context. Non-spatial BTSP described in this paper and spatial BTSP described in spatial contexts [3–12] share many attributes. First, they are both induced by large calcium events. Second, like spatial BTSP, odor-responsive fields typically form around 0.5 seconds before the time of onset of the ‘plateau-like’ event. This temporally asymmetric induction of fields is characteristic of BTSP in CA1. Membrane potential (Vm) recordings in CA1 during spatial BTSP demonstrate potentiation causing the induction of an asymmetric Vm ramp extending back nearly 4 seconds from the timepoint of induction. We infer potentiation by visualizing asymmetric firing rate increase ramps that are remarkably similar to those Vm ramps recorded in spatial tasks; however, voltage recordings would be required to determine whether a Vm ramp extending several seconds is also induced by non-spatial BTSP. Third, we can induce BTSP-like formation of odor responses with a single large stimulation of one CA1 pyramidal neuron. Fourth, we find that rates of BTSP and potentiation are highest during the first day of learning. This corroborates the findings that BTSP is a neural mechanism underlying learning of tasks and novelty.

There are notable differences between spatial and non-spatial BTSP, however. While spatial BTSP can induce place fields anywhere in the virtual track, during our non-spatial task, 86% of successful BTSP-like fields were formed during or immediately after the odor presentations, with few fields formed during the delay and reward periods. The firing rate increase potentiation for plateau-like events during the delay period were dramatically lower than those during the odor presentation, and we were unable to induce any fields during the delay period with holographic optogenetic stimulation. It is possible that this occurs because subthreshold inputs potentiated by BTSP in the delay period fail to reach action potential threshold. Perhaps several stronger induction events are necessary to form representations during the delay period. This could be explained by the fewer and weaker EC inputs activated during the delay period as LEC has nearly twice as many axons and MEC nearly 10 times as many axons with peak firing during the odor period compared to the delay period. Recordings of Vm during the task would be necessary to find whether the magnitude of synaptic potentiation is similar during the different phases of the task. While our findings of MEC inhibition reducing plateau-like event rates corroborate findings that MEC TA inputs drive plateau potentials in spatial tasks [9], it remains to be determined whether TA inputs, CA3 inputs, or both are potentiated during non-spatial BTSP. Finally, while inhibitory interneuron subtypes have been characterized by their roles in gating spatial EC and CA3 inputs to CA1 [47–50], it remains unclear how the different interneuron subtypes within the different layers of CA1 contribute to BTSP and gate non-spatial sensory inputs. Future recordings and manipulations of the activity of these neurons can further elucidate the complex mechanisms underlying non-spatial BTSP in CA1.

We also find that inhibition of LEC and MEC have distinct differential effects on non-spatial BTSP. While MEC inhibition reduces the frequency of large calcium events, LEC inhibition had no impact on the frequency or amplitude of these events but reduced their potentiation and success rate in generating odor-fields. This indicates that MEC plays a major role in generating the plateau potential teaching signal with most of its activity timed to stimulus presentations. In contrast, the exact mechanism through which LEC regulates the success of BTSP events is less clear. There are several possibilities. It is possible that BTSP potentiates the LEC inputs on the distal dendrites of CA1 pyramidal neuron which aids in generating odor-selective responses. More likely, it is possible that LEC inhibition reduces odor-selectivity and the amplitude of odor responses in dentate gyrus granule neurons or in CA3, which in turn reduces the potentiation of CA3 inputs to CA1. Our results are in line with studies which have shown the importance of MEC inputs for generation of teaching signals to drive large responses in CA1 and BTSP during spatial learning tasks [7, 9, 51], while indicating further complexity given the distinct roles of LEC and MEC. Our findings support the structural and functional connectivity of LEC and the hippocampus in olfactory based tasks [32–35], but further experiments with other modalities would be valuable in establishing LEC and MEC’s unique roles in driving plateau potentials and forming non-spatial representational fields. CA1 is also well known for its internal representations [21–23]. Although we observed small firing rate increases and rare BTSP-like events that form odor-specific fields during the delay period, future recordings should investigate if LEC and MEC inputs coincide with the output from recurrent CA3 networks capable of generating temporal codes [52–54] to drive BTSP for internally generated representations.

Despite research of CA1 dendritic activity during learning of spatial environments [11, 55–57], the molecular mechanisms underlying BTSP are still poorly understood. While CaMKII has been demonstrated to be critical for BTSP [10, 58], the role of neuromodulators like acetylcholine and norepinephrine are less understood [59]. Finally, while inhibitory interneuron subtypes have been characterized by their roles in gating spatial EC and CA3 inputs to CA1 [47–50], questions remain to how the different interneuron subtypes within the different layers of CA1 contribute to BTSP [60] and gate non-spatial sensory inputs. Future recordings and manipulations to investigate the role of neuromodulators and activity of dendrites and inhibitory neurons will further elucidate the complex mechanisms underlying non-spatial BTSP in CA1.

## ACKNOWLEDGMENTS

We thank D. Buonomano, G. Orban, and S. Miller-Hecht for feedback on the manuscript. We also thank B. Madruga, Z. Day, X. Chen, and C. Yang for technical support. This work was supported by the following funding sources: T32DGE1829071, 5T32NS045540-20, R01NS116589, and 1P50HD103557-01 (P.G.). Whitehall Foundation Grant and K08NS109315 (A.A.).

## AUTHOR CONTRIBUTIONS

C.D., J.T., and P.G. conceived the experiments. C.D. and A.A. conducted holographic optogenetics experiments. C.D. conducted all other experiments, analyzed experimental data, and generated the figures. C.D. and P.G. wrote the manuscript.

## RESOURCE AVAILABILITY

### Lead Contact

Further information and requests for resources and reagents should be directed to the Lead Contact, Peyman Golshani.

### Material Availability

No new materials were created for this study.

### Data and Code Availability

Data and analysis code will be made available on Dryad and GitHub after manuscript acceptance. Any additional information necessary to reanalyze the data reported in this paper is available from the lead contacts.

## EXPERIMENTAL MODEL AND SUBJECT DETAILS

### Animals

All of the experiments were conducted according to the National Institute of Health (NIH) guidelines and with the approval of the Chancellor’s Animal Research Committee of the University of California, Los Angeles. A total of 9 adult male and 8 female mice (8-16 weeks old, C57BL/6J The Jackson Laboratory 000664) were used for simultaneous *in vivo* calcium CA1 imaging and EC chemogenetic experiments (Figures 1 and 4). These mice were divided into 4 groups: LEC mCherry n=3, MEC mCherry n=3, LEC PSAM4 n=6, MEC PSAM4 n=5. A total of 3 adult female mice (25-35 weeks old) were used for simultaneous *in vivo* calcium CA1 imaging and holographic optogenetic stimulation experiments (Figure 2). These 3 positive offspring were chosen from a breeding of B6;DBA-Tg(tetO-GCaMP6s)2Niell/J (The Jackson Laboratory 024742) or B6;D2-Tg(tetO-GCaMP8s)1Genie/J (The Jackson Laboratory 037717) crossed with B6.Cg-Tg(Camk2a-tTA)1Mmay/DboJ (The Jackson Laboratory 007004). A total of 2 adult male and 3 female mice (8-16 weeks old, C57BL/6J The Jackson Laboratory 000664) were used for *in vivo* calcium CA1 imaging during learning experiments (Figure 3). A total of 7 adult male and 9 female mice (8-16 weeks old, C57BL/6J The Jackson Laboratory 000664) were used for *in vivo* calcium EC axon imaging experiments (Figure 5). These mice were divided into 2 groups: LEC n=8, MEC n=8. All were experimentally naïve and housed in the vivarium under a 12-hour light/dark cycle. All mice were group housed (2-4 per cage) except for 2 that had to be separated following surgery because of fighting.

## METHOD DETAILS

### Surgical Procedures

Mice were subcutaneously administered pre-operative drugs (carprofen 5 mg/kg, dexamethasone 0.2 mg/kg, lidocaine 5 mg/kg) 30 minutes before surgery. Mice were anaesthetized with isoflurane (5% induction, 1-2% for maintenance), and anesthesia was continuously monitored and adjusted as necessary. The scalp was shaved, and mice were placed into a stereotactic frame (David Kopf Instruments, Tujunga, CA) on a feedback-controlled heating pad (Harvard Apparatus) set to maintain body temperature at 37*^◦^*C. Eyes were protected from desiccation using artificial tear ointment. The surgical incision site was cleaned three times with 10% povidone-iodine and 70% ethanol. Fascia was removed by applying hydrogen peroxide, connective tissue was cleared from the skull, and the skull was scored to facilitate effective bonding with adhesives at the end of surgery. After stereotactically aligning the skull, a single or several burr holes were made depending on the experiment performed and virus was injected.

CA1 imaging with EC chemogenetics experiments: Control virus (500 nL of 1:5 saline dilution of pAAV1-CaMKIIa-mCherry into all 4 sites) or experimental virus (500 nL of 1:5 saline dilution of AAV5-CaMKII-PSAM4-GlyR-IRES-EGFP into all 4 sites) was injected into LEC (bilaterally 3.4 and 3.9 mm posterior, 4.35 mm lateral, and 4.3 ventral from bregma) or MEC (bilaterally 4.7 mm posterior, 3.35 mm lateral, and 3.8 and 3.0 mm ventral from bregma). Additionally, pGP-AAV1-syn-jGCaMP8f-WPRE (1000nL of 1:5 saline dilution) was injected into the right dorsal CA1 (2.0 mm posterior from bregma, 1.8 lateral from bregma, and 1.3 ventral from dura).

CA1 imaging with holographic optogenetics experiments: AAV8-CaMKIIa-ChRmine-mScarlet-Kv2.1-WPRE (1000nL of 1:10 saline dilution) was injected into the right dorsal CA1 (same coordinates as above).

CA1 imaging during learning experiments: pGP-AAV1-syn-jGCaMP8f-WPRE (1000nL of 1:5 saline dilution) was injected into the right dorsal CA1 (same coordinates as above).

EC axon imaging experiments: pENN.AAV1.CaMKII.0.4.Cre.SV40 and pGP-AAV1-CAG-FLEX-jGCaMP7f-WPRE were mixed immediately before the injection (500 nL of 1:1 mix) into right LEC (3.5 mm posterior, 4.35 mm lateral, and 4.3 ventral from bregma) or right MEC (4.7 mm posterior, 3.35 mm lateral, and 3.5 mm ventral from bregma). All viruses were injected using a Nanoject II microinjector (Drummond Scientific) at 60nL per minute.

For mice in all experiments, following virus injection, a circular craniotomy (3 mm diameter) was made centered around a point made 2.0 mm posterior and 1.8 lateral to bregma. Dura beneath the craniotomy was removed and cortical tissue above dorsal CA1 was carefully aspirated using a 27-gauge blunt needle. Corpus callosum was spread to the sides of the craniotomy to expose the alveus. Cortex buffer (NaCl = 7.88g/L, KCl = 0.372g/L, HEPES = 1.192g/L, CaCl_2_ = 0.264g/L, MgCl_2_ = 0.204g/L, at a pH of 7.4) was continuously flushed during aspiration and until bleeding stopped. A titanium ring with a 3 mm diameter circular thin #0 coverglass attached to its bottom was implanted into the aspirated craniotomy and the overhanging flange was secured to the skull with vetbond (3M). A custom-made lightweight stainless-steel headbar was attached to posterior skull and secured with cyanoacrylate glue. Dental cement (Ortho-Jet, Lang Dental) was applied to seal and cover any remaining skull, and to form a small well around the titanium ring for holding immersion water for the objective during imaging. Following surgery, all animals were given post-operative care (carprofen 5 mg/kg and dexamethasone 0.2 mg/kg for 48 hours after surgery) and provided amoxicillin-treated water at 0.5 mg/mL for 7 days. All mice recovered for 7-14 days before experiments began.

### Experimental setup

The entire behavioral setup is as described in Taxidis et al. [20]. Mice were head-fixed above an 8-inch spherical Styrofoam ball (Graham Sweet) which can rotate about one axis for 1D locomotion that was recorded with a sensor (Avago ADNS-9500). A continuous stream of clean air (∼1 L/min) was delivered toward the animal’s nose via Tygon PVC clear tubing and a custom-made port that held the air tube and water port. At the onset of the odor presentation period, a dual synchronous 3-way valve (NResearch) switched to the odorized one for 1 second. Odorized air was created by using a 4-ports olfactometer (Rev. 7c; Biology Electronics, Caltech) supplying air to either of two glass vials containing odor A (70% isoamyl acetete basis, FCC; Sigma Aldrich) or odor B ((−)-a-Pinene ≥ 97%, FCC; Sigma Aldrich), which were both diluted in mineral oil at 5% concentration. Water droplets (∼10µl) were released by a 3-way solenoid valve (Lee Company), and licks were detected by using a custom battery-operated circuit board with one end of the circuit connected to the headbar and the other to the lickport. The behavioral rig was controlled with custom written software (MATLAB) and through a data acquisition board (USB-6341: National Instruments).

### Behavioral training

After 7-14 days recovering from surgery, mice were handled and began water-restriction to 85% of their original weight before water-restriction. After one day of handling, mice were habituated to being head-fixed above the spherical treadmill for two days.

All experiments except CA1 imaging during learning: On the 4th day of training, mice began learning to lick from the lickport as water was automatically delivered at the beginning of the reward period following only non-matched odor trials (AB or BA, with water delivery at time point of 8 seconds). This phase was always 2 days except for the rare mouse that needed one extra day to reach motivation level and lick water from the port for at least 50 trials. In the next phase, water was only delivered if the mouse licked during the response period, and mice learned to reliably lick in anticipation of the reward following the 2^nd^ odor. This phase was also 2 or 3 days, dependent on the mouse licking during the response period of at least 50 trials. The final phase was the full delayed non-match-to-sample (DNMS) task in which matched odor trials (AA and BB) were introduced and mice learned to refrain from licking the port following these trials.

CA1 imaging during learning experiments: All previous training steps were identical, except that the olfactometer remained off until Day 1 (which was the first day that mice received any air or odors). Two-photon calcium imaging was conducted on the last 2 days when water was only delivered if the mouse licked during the response period (baseline days referred to as Day −2 and −1 in the text and figures). For all 11 imaging days, a total of 100 trials delivered in five blocks of 20 trials were given each day. All 9 imaging days across learning of the DNMS task were conducted consecutively and began within 3 hours of the start time of the previous day.

CA1 imaging with EC chemogenetics experiments: Mice underwent 5-7 days of learning the full DNMS task before recording began. Two-photon calcium imaging only began after the mouse had 2 consecutive days of ‘expert performance’ (at least 85%). For all imaging days, a total of 100 trials delivered in five blocks of 20 trials were given each day. All 8 imaging days were conducted consecutively and began within 3 hours of the start time of the previous day.

CA1 imaging with holographic optogenetics experiments: Mice underwent at least 7 days of learning the full DNMS task before recording began. For all imaging days, 60 trials (three blocks of 20 trials) were given for each FOV (up to 3 FOV sessions per day). Imaging days were not consecutive because we did not match FOVs and performance was stable.

EC axon imaging experiments: Imaging was conducted for 7 days and began before expert performance of the DNMS task. However, only sessions with expert performance were included in analysis. All 7 imaging days were conducted consecutively and began within 3 hours of the start time of the previous day.

All experiments: Trials were delivered in blocks of 20 trials. There was no punishment or timeout following an incorrect lick; the water was simply not delivered.

### *In vivo* two-photon imaging

All two-photon calcium imaging without holographic optogenetics was conducted using a resonant scanning two-photon microscope (Scientifica) fitted with a 16x 0.80 NA objective (Nikon) to record 512×512 pixel frames at 30.9 Hz. CA1 imaging fields of view were 500×500 µm and axonal imaging fields were 250×250 µm. Excitation light was delivered with a Ti:sapphire excitation laser (Chameleon Ultra II, Coherent), operated at 920 nm. GCaMP8f and GCaMP7s fluorescence was recorded with a green channel gallium arsenide photomultiplier tube (GaAsP PMT; Hamamatsu). Microscope control and image acquisition were performed using LabView-based software (SciScan). Imaging and behavioral data were synchronized by recording TTL pulses generated at the onset of each imaging frame and olfactory stimulation digital signals at 1 kHz, using WinEDR software (Strathclyde Electrophysiology Software).

CA1 imaging experiments: A single field of view (FOV) was imaged for all recordings. Careful attention was given to aligning the FOV to the previous day’s as perfectly as possible. Animals were not included in analysis if successful alignment was not possible. We used rotating stages, a motor for adjusting mouse head angle, and a tiltable objective attachment with two degrees of freedom to fine-tune the alignment.

EC axon imaging experiments: The same alignment was always attempted for 7 consecutive days, but the extra difficulty of alignment made it not always possible. Therefore, axon segments were not registered between days; however, FOVs were typically very similar. Laser power and PMT settings were kept consistent between days, except for rare occasions when it was necessary to maintain similar signal-to-noise. Out of the 16 axonal imaging animals included in analysis (each recorded for 7 days), 42 recording sessions were not included because of sub-expert performance and 7 sessions were not included because of poor signal-to-noise (resulting in a total of 63 included recording sessions across the 16 animals).

For each day of recording, imaging was halted between each of the 5 blocks of 20 trials. This allowed fine-tuning of alignment, and it also prevented brain heating or photo-toxicity. Laser power was kept as minimal as possible (60-80mW for CA1, and 100-200mW for EC axons) without sacrificing signal-to-noise ratio, and only mild photo-bleaching was observed in some axonal imaging animals.

### *In vivo* two-photon imaging with holographic optogenetic stimulation

Two-photon calcium imaging with holographic optogenetics was conducted using a resonant scanning two-photon microscope (Ultima 2Pplus, Bruker) fitted with a 16x 0.8 NA objective (Nikon) to record 512×512 pixel frames at 30 Hz. CA1 imaging FOVs were 556×556 µm. GCaMP excitation light was delivered with an excitation laser (InSight X3, SpectraPhysics-Newport) operated at 920 nm. GCaMP6s and GCaMP8s fluorescence was recorded with a green channel gallium arsenide photomultiplier tube (GaAsP PMT; Hamamatsu). ChRmine excitation light was delivered with another excitation laser (Spirit One 1040-8, SpectraPhysics-Newport) operated at 1040 nm. Timing of stimulation was triggered from the behavioral setup that also triggered odor and water delivery.

After locating a suitable FOV, individual neurons were tested for ChRmine excitation by delivering one or two small stimulations (five 10ms pulses at 10Hz). Laser power for each stimulation was kept at a minimum level that would induce stimulation. If the neuron responded with a noticeable increase in fluorescence, a point was placed. This was repeated until up to 8 neurons had been selected in the ROI manager. Then behavior was started and stimulation points were delivered one-by-one starting on the 5^th^ trial. These large stimulations (ten 10ms pulses at 10Hz lasting one second) were meant to induce large-amplitude, long-duration calcium events resembling plateau potentials. The timing of stimulation was predetermined with a higher concentration near the offset of odors and an even distribution through the delay and reward periods. Another different 8 neurons were selected for the next block of 20 trials for the FOV, and the steps were repeated. With up to 3 sessions (60 trials) per FOV, up to 24 neurons were stimulated. If additional FOVs were suitable, the previous steps were repeated, and up to 3 FOVs were imaged in a single day (for up to 72 neurons total across 3 sessions of 60 trials each).

Mice were given at least one rest day between recordings, and a maximum of 3 days were conducted per animal.

### Chemogenetic inhibition

All CA1 imaging with EC chemogenetics animals received subcutaneous injections of saline for at least 5 days prior to imaging to habituate them to the injection prior to being head-fixed. For the 8 days of imaging, mice received alternating injections of saline and uPSEM (ultrapotent PSEM 792 hydrochloride binds to PSAM4 to cause strong inhibition). Half of the mice started with saline and the other half started with uPSEM on the first day of imaging. The uPSEM powder was dissolved into saline at a concentration of 0.3 mg/mL, and injections were administered to achieve 3 mg/kg. After weighing the mouse to calculate the appropriate volume of saline or uPSEM, the mouse was injected intraperitoneally and head-fixed under the microscope. 10-20 minutes elapsed between the injection and the start of behavior and imaging.

### Histology

Following all experiments, mice were deeply anaesthetized under isoflurane and transcardially perfused with 30 mL 1x PBS followed by 30 mL 4% paraformaldehyde in 1x PBS at a rate of approximately 4 mL/min. After perfusion, the brains were extracted and post-fixed in 4% paraformaldehyde. Sections of 80 µm were collected using a vibratome, 24-48 hours after perfusion. For animals with LEC viral expression, coronal sections were taken, while sagittal sections were taken from animals with MEC viral expression. The sections were mounted onto glass slides and cover-slipped with DAPI mounting medium. Images were acquired on an Apotome2 microscope (Zeiss; 5x, 10x, 20x objectives) to confirm proper expression and location of viral expression. For chemogenetic experiments, sufficient PSAM4 or mCherry expression was found restricted to either LEC or MEC. In axonal imaging experiments, somatic GCaMP7s was confirmed to be restricted to only LEC or MEC, and axonal expression was found in the SLM layer of dorsal hippocampus. Mice with insufficient PSAM4/mCherry expression or PSAM4/mCherry/GCaMP7s that spread outside of their desired target were excluded from analysis.

## QUANTIFICATION AND STATISTICAL ANALYSIS

### Calcium imaging data pre-processing

CA1 imaging with EC chemogenetics experiments: The 8 days of recordings were divided into 4 pairs of days, so that each pair consisted of one saline day and one uPSEM day. Both recordings from a single pair were concatenated before processing so that the same neurons could be detected within the pair of imaging days. Concatenated movies were processed using the Python implementation of Suite2P 0.9.2 [41] to perform non-rigid motion registration, neuron segmentation, extraction of fluorescence signals, and deconvolution with parameters optimized to our GCaMP8f CA1 recordings. We used the default classifier and an ‘iscell’ threshold of 0.1 to only include masks that were likely neurons.

CA1 imaging during learning experiments: All parameters were the same as previous, but days were processed separately.

CA1 imaging with holographic experiments: Each FOV was processed separately. GCaMP6s and GCaMP8s recordings had the same parameters are the previous sections except for ‘tau’ (timescale of sensor), was set to 1.25 and 0.5, instead of 0.25. Stimulation ROIs were manually verified. During some stimulations, more than one neuron responded to a single stimulation. This was due to brain movement and the high density of CA1 neurons. Any neighboring neuron that also responded to the stimulation was included as a stimulated neuron. Some neurons failed to respond at all (left bin in Fig. 2d).

EC axon imaging experiments: The 7 days of recordings were all processed separately. Movies were also processed using Suite2P but with parameters optimized to our GCaMP7s axonal recordings. An additional step of axon merging was taken to decrease the number of duplicates (as an axon could appear as multiple segments within the FOV); this also increased signal-to-noise by increasing the number of pixels for a single mask. By visualizing axon correlation values and their fluorescence traces within the Suite2P GUI, we chose axon segments to merge based on correlation values and footprint distributions. Using custom Python code with functions from Suite2P’s source code, we ‘merged’ axons by generating new ROIs with these new pixels. The old axon segments were then eliminated from analysis and deconvolution was run on the new axon masks.

For all experiments, deconvolved signals were taken as the selected output from Suite2P and analyzed further in MATLAB 2021a. Deconvolved signals were smoothed by a rolling mean of 10 frames (0.32 seconds), then z-scored, and finally values below 2 were set to zero. The resulting signal was used for all analysis and referred to as ‘firing rate (STD)’ as a proxy for spiking activity. Signals were aligned to the trial structure (odor presentations, reward period, lick timing) and the recorded locomotion as mice ran on the spherical ball.

### Binary BTSP event detection and analysis

First, 6-second periods were extracted for each odor presentation period (2 seconds before odor and 3 seconds after) and divided for Odor A and Odor B regardless of whether it was the first or second odor presented in the trial. Since each recording had 5 blocks of 20 trials, we have 100 odor presentations of each odor per neuron per day. Next, we identified every single calcium event, which we defined as a group of consecutive timepoints with a non-zero deconvolved signal. The size and timing of that event are counted as the peak value within the event and that timepoint’s time relative to the odor, respectively.

Next, we identified which events satisfied criteria to be considered as a possible induction event. This detection was performed separately for Odor A and Odor B presentations. Events in the first 10 or last 10 odor presentations were not considered for analysis, because we needed enough odor presentations before and after the event to detect BTSP events. There were two criteria for an event to be considered a possible induction event: during the previous 3 odor presentations the neuron must show no activity within 2 seconds before or after the event in question, and there must not have been a significant peak firing field. To determine the significance of a firing field, we took 6-second periods of all previous odor presentations and found the peak of the average activity. We then circularly shuffled each odor presentation and found the peak of the average activity from this shuffled data. This was repeated 2000 times to generate 2000 peak values from shuffled data. For a possible induction event, the real peak of average activity must not have been greater than the 90^th^ percentile of the shuffle.

If an event passed criteria to be considered as a possible induction event, we analyzed if it was successful in forming a field. There were four criteria for a successful field formation: 1. The resulting field must have been significant above the 95^th^ percentile of the shuffle; 2. The resulting field occurred within 2 seconds of the peak of the induction event; 3. The neuron must have fired (have value above 2 STD) within 0.5 seconds of the resulting field for the next 3 odor presentations; 4. The neuron must have fired within 0.5 seconds of the resulting field for at least 7 out of the next 10 odor presentations. The strict criteria for activity in the previous 3, following 3, and following 10 odor presentations improves the likelihood that the event in question does induce a stable field. The ± 2 second window for the difference between the event peak and field peak allowed us to look for asymmetrical formation without any bias. The lack of any criteria regarding the amplitude of the induction event allowed us to probe the relationship of amplitude to success rate and asymmetrical formation. Success rate increased continuously with amplitude (Fig. 1i), but only events with amplitude above 10 STD had statistically significant asymmetrical formation. Therefore, we considered any reference event above 10 STD to be ‘plateau-like’, and successful ‘plateau-like’ events are what we considered to be ‘BTSP-like’ events. We considered any event between 2 and 10 STD to be a ‘small event’.

### Firing rate increase analysis

As in the previous section, every single calcium event was identified. Events were not included in analysis if there were not at least 10 trials of the same and 10 trials of the opposite trial type both before and after the trial of the reference event. For the ‘same odor’ analysis, the previous 10 trials of the same odor type were averaged and subtracted from the average of the following 10 trials of the same odor type. The same was done for ‘opposite odor’ analysis but with the trials of the opposite trial type. The firing rate increase ramps were aligned to the peak of the reference event and averaged for all references of the desired specifications. Displayed ramps are not binned and show datapoints from each recorded frame.

When the firing rate increase value is collected as a single value (not displaying ramp), the 30 frames of the 1 second period from −1 to 0 (aligned to reference event) were averaged. This value was then averaged for all events within a single recording session.

For figures showing the ramps or firing rate increases across the trial structure, the entire trial structure was used for the calculation. For figures showing the effect only during the odor period, odor presentations were isolated as done in the previous section (ignoring whether the odor came first or second).

### Locomotion analysis

1D locomotion that was recorded with a sensor (Avago ADNS-9500) at 1kHz was binned to match the frame rate of calcium imaging. The signals were binned to 1/6 seconds, z-scored, and finally values between −1 and 1 were set to 0. Any bin with a locomotion value of 0 was determined to a period of ‘no movement’. A bin or group of bins with values not equal to 0 and lasting for less than one second were considered ‘minor jitter movement’. This included positive and negative values and the minor jitter movements included small ball movements in both directions. A group of bins with values less than −1 and lasting for at least one second were considered bout of ‘running’. The negative is indicative of the ball rolling backwards (which is caused by the mouse running forward).

### Selectivity analysis

We calculated the odor-selectivity index value for each ROI as: SI = (R_a_ - R_b_) / (R_a_ + R_b_); where R_a_ is the firing rate at a given bin for Odor A trials and R_b_ is the same for Odor B trials. Bin sizes were always 0.5 seconds, and all trials were included. For each ROI, a distribution of 2000 shuffled index values were also calculated by randomly shuffling the trial type assignment 2000 times for each bin. The maximal absolute value index is chosen from all the bins (for the real ROI and all 2000 shuffles), and the bin is noted. ROI’s with an absolute value index value above the 95^th^ percentile of absolute value shuffled index values are considered to be ‘significantly selective’. For figures displaying the actual odor-selectivity index value (ranging from 0 to 1), the values of the 2 bins during odor presentations were simply averaged.

### Support vector machine decoding

Binary support vector machine (SVM) decoding was performed in MATLAB 2021a (default parameters) using bin sizes of 0.5 seconds (averaging the deconvolved signal for those frames within the bin).

CA1 imaging with EC chemogenetics experiments: The number of ROIs was controlled by randomly subsampling 100 ROIs out of all possible ROIs. The result of 20 subsamples of ROIs were averaged for each data point. For each bin and subsample, 80% of trials were used for training the decoder, and the remaining 20% were used for testing. This was repeated 4 more times so that each block of 20 trials was used as the 20% for testing. For each training of the decoder, another training was done with a shuffled assignment of trial type to confirm a shuffle comparison of data yields decoder accuracy of ∼50%. For odor decoding, the trials were broken down into odor presentations (same as in BTSP detection analysis) to evaluate odor decoder accuracy regardless of the order of the odors. When specific timepoints were mentioned, such as ‘during odor presentation’ or ‘during delay period’, the average accuracies of the 0.5 second bins were averaged (not trained/tested with larger bins).

EC axon imaging experiments: To control the number of axon segments, the 10 most odor-selective neurons were selected from the previous analysis (those with the highest percentile values). All other parameters remained the same.

### Sequence-axon detection and analysis

To evaluate peak firing timing in EC axons, we performed sequence-axon detection similar to the previously described approach in CA1 neurons in our DNMS task, Taxidis et al. [20]. First, trials that begin with Odor A and those that begin with Odor B were separated, and the one with a larger peak of the average activity was considered further. Additionally, only the 6-second period including first odor presentation and the delay period was considered. In the same way as described in BTSP-event detection, the peak of average activity within this period and a given trial type was determined to be significant if the peak was greater than the 95^th^ percentile of 2000 circular shuffles. The neuron must also have had a trial reliability of at least 20% (have fired above 2 STD for 20% of the preferred trials within 0.5 seconds of the peak frame found in the previous step). If an ROI passed both criteria, it was considered to be a ‘sequence-axon’ regardless of its odor-selectivity, as that was a separate analysis. An ROI was considered an ‘odor axon’ if its peak was within the odor presentation period. An ROI was considered an ‘offset axon’ if its peak was in the first second of the delay period. An ROI was considered a ‘delay axon’ if its peak was during the last 4 seconds of the seconds of the delay. This was done to not include the large population of ROIs that fired to the offset of the odor (likely the auditory cue of the clicking of the valve).

### Statistical analysis

All statistics were performed in MATLAB 2021a and described in figure legends. In all cases in the text, values were written in the format ‘mean ± standard deviation’ (STD), while error bars in all figures show the mean and standard error of the mean (SEM). For all figures, no asterisks were shown if p ≥ 0.05, 1 asterisk if p < 0.05, 2 asterisks if p < 0.01, 3 asterisks if p < 0.001. If the p ≥ 0.1, ‘n.s.’ is displayed, but if 0.05 ≤ p < 0.1 the p-value was typically displayed in the figure. On occasions when single asterisks were displayed above a curve or trace, p-values were corrected for multiple comparisons using the false discovery rate Benjamini-Hochberg procedure. No statistical methods were used to determine appropriate sample sizes but were chosen as being comparable to sizes used in similar publications.

## SUPPLEMENTAL FIGURES S1-S8

**Supplemental Figure 1:**
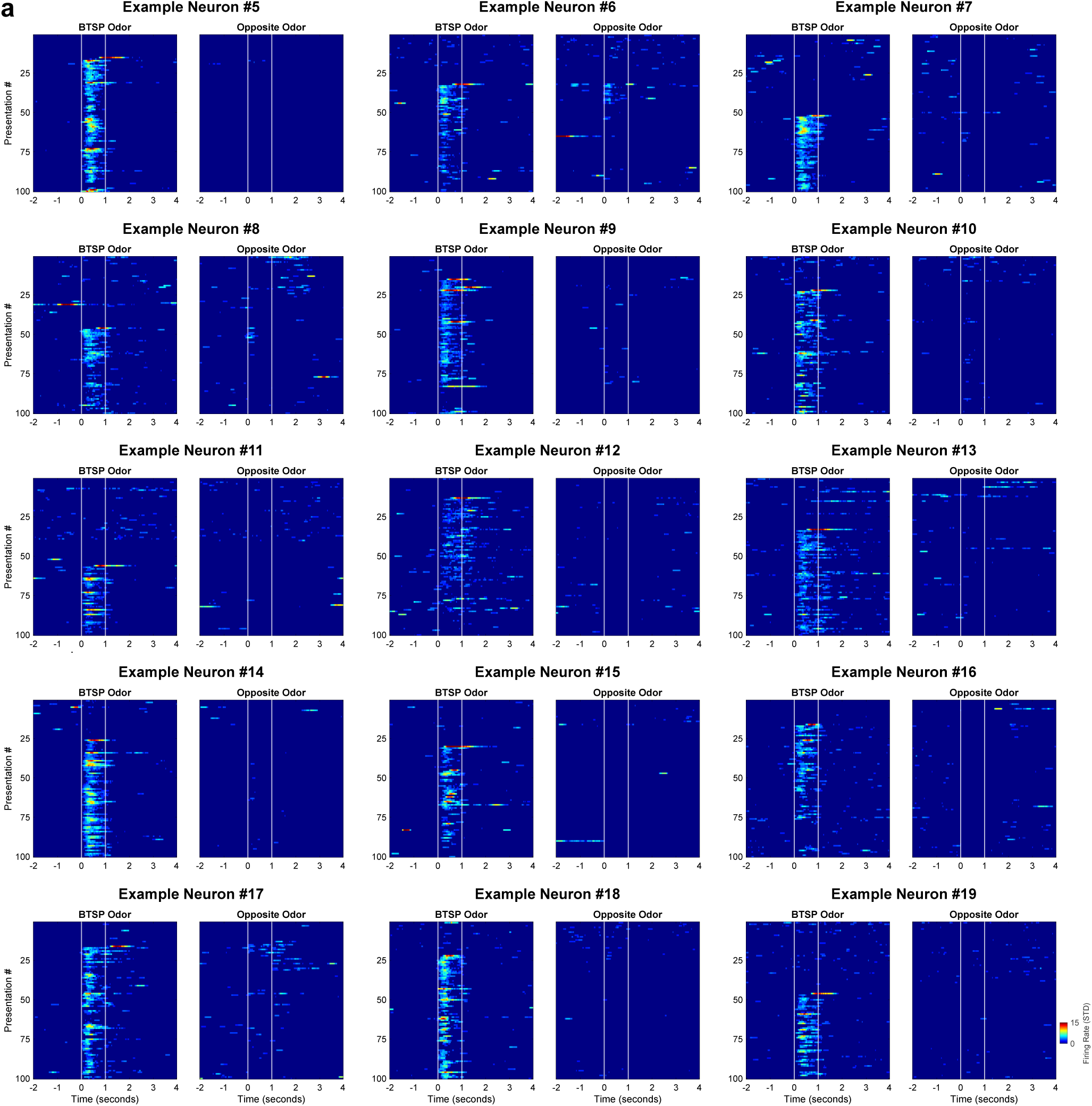
Additional examples of ‘BTSP-like’ events. **a,** 15 more example neurons showing ‘BTSP-like’ events.

**Supplemental Figure 2:**
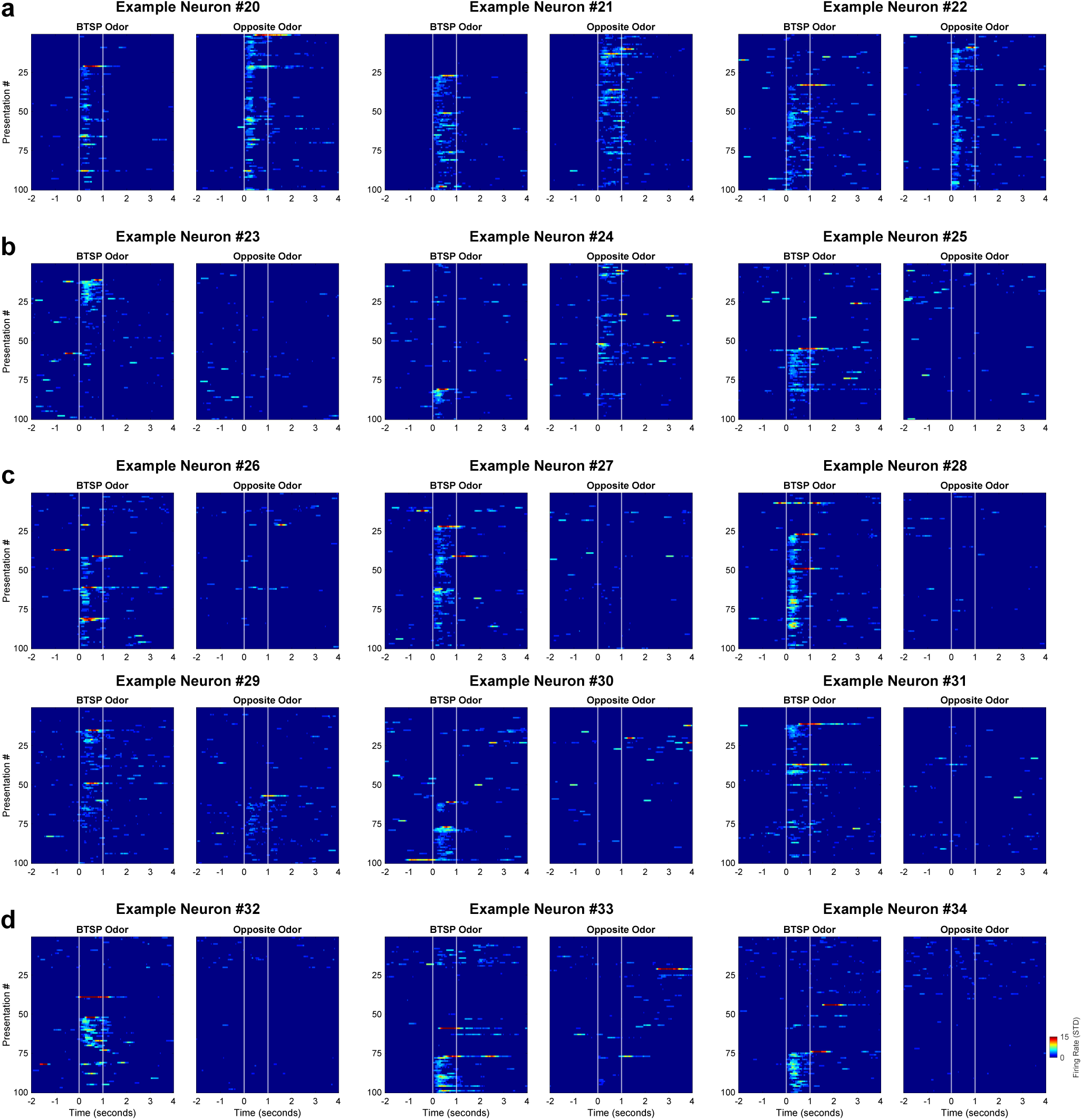
More unique examples of ‘BTSP-like’ events. **a,** 3 examples of an odor-selective neuron becoming non-selective because of the new BTSP induced field. **b,** 3 examples of neurons that formed an odor-field after a ‘plateau-like’ event, but the field faded quickly. **c,** 6 examples of neurons with multiple ‘plateau-like’ events that seem to reinforce the odor-field. **d,** 3 examples of failed ‘plateau-like’ events followed by successful ones.

**Supplemental Figure 3:**
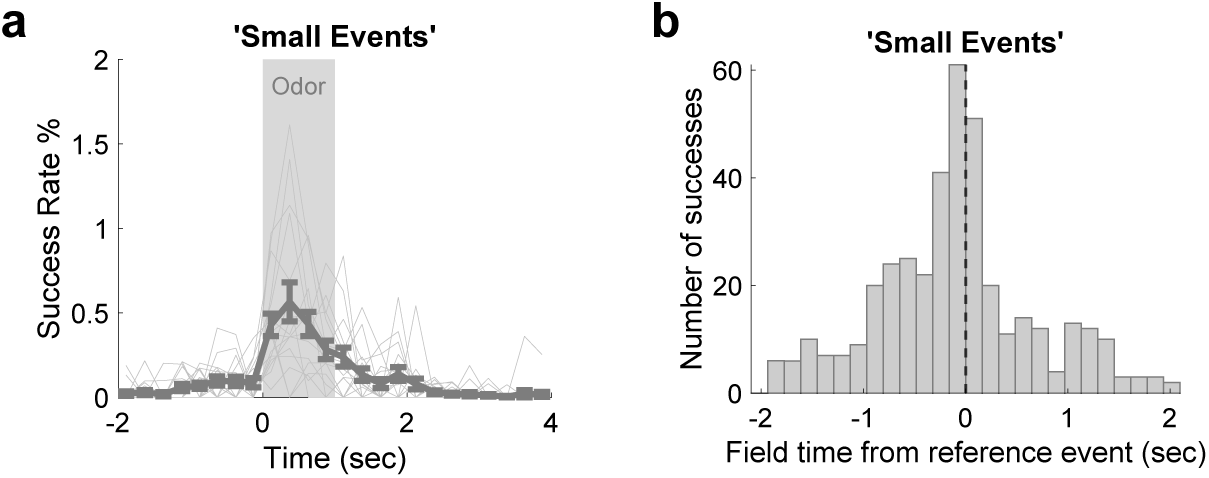
Small events were not BTSP. **a,** Supplementing Fig. 1j showing success rate was very low for small events. These represent randomness of non-BTSP events passing criteria for BTSP. **b,** Supplementing Fig. 1k showing that small events did not have asymmetrical formation that was seen for ‘plateau-like’ events (n=396).

**Supplemental Figure 4:**
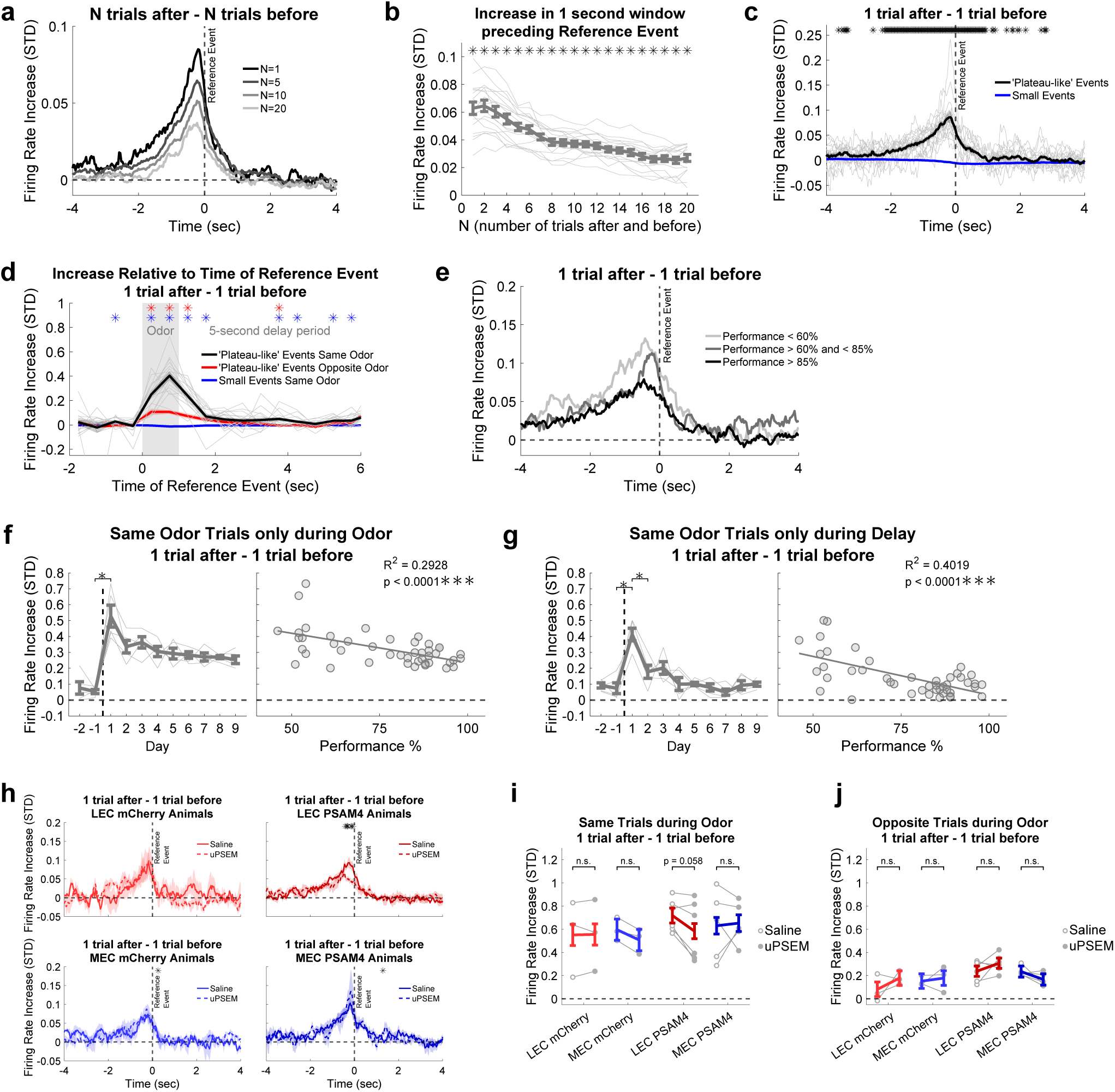
Firing rate increase effects were found at a range of number of trials after and before. **a,** Supplementing Fig. 1l to show the firing rate increase ramp with 4 different number of trials (N) after and before. **b,** Calculated as the average of the ramp between −1 and 0 seconds in trial-time, plotted as a function of N. Statistics are one-sample t-tests with each N having a p < 0.05 (corrected for multiple comparisons using the Benjamini-Hochberg procedure). N = 10 was chosen for the main figures because it minimized variability. N = 1 has the largest effect but largest variability. **c,** Same as Fig. 1l but for only 1 trial after and 1 trial before. **d,** Same as Fig. 1n. **e,** Same as Fig. 3f. **f,** Same as Fig. 3g. **g,** Same as Fig. 3i. **h,** Same as Fig. 4d. **i,** Same as Fig. 4e. **j,** Same as Fig. 4f.

**Supplemental Figure 5:**
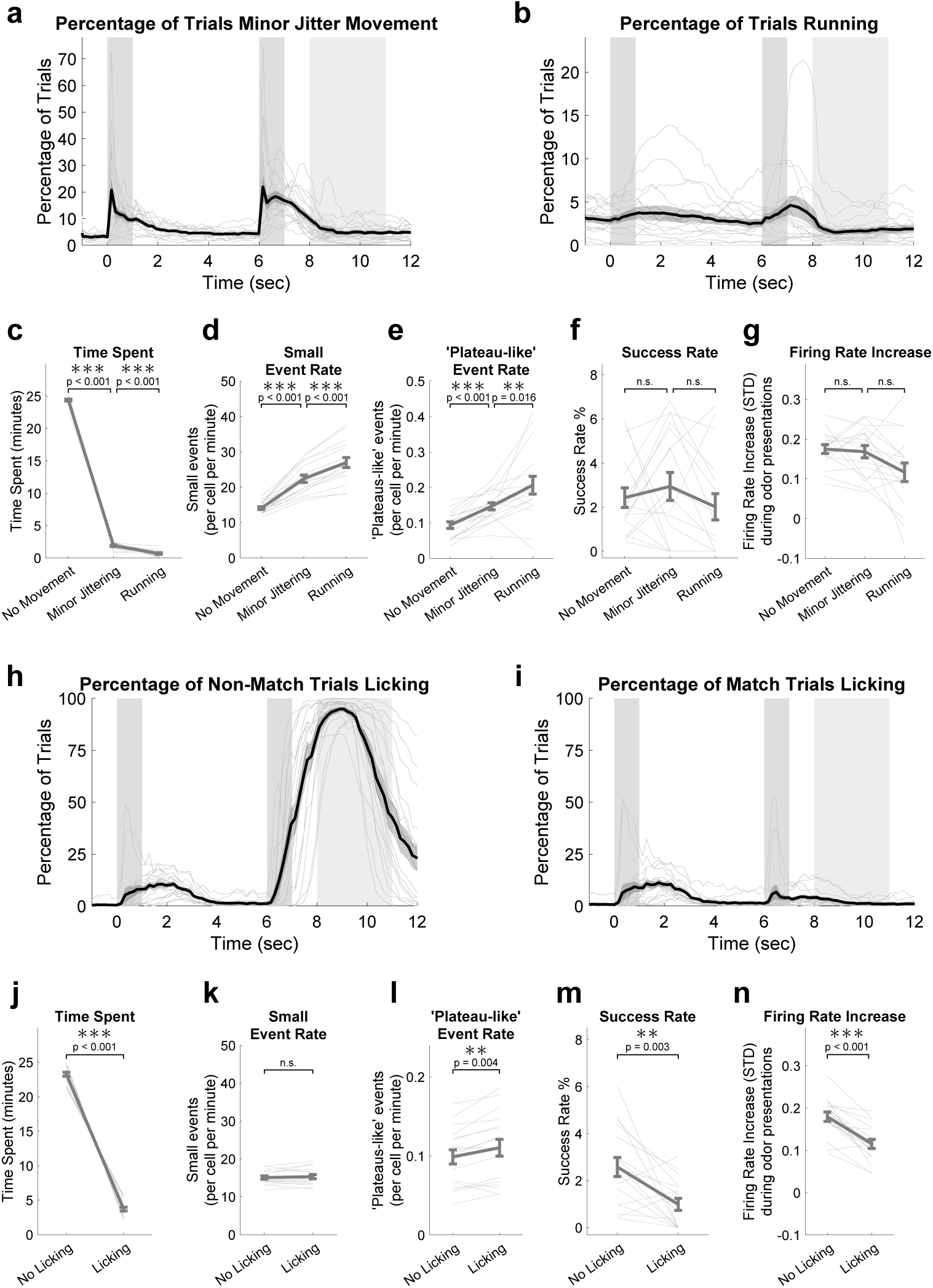
All types of movements increased neural activity but had no effect on BTSP. **a,** Percentage of trials with a minor jitter movement at each bin in the trial. Minor jitter movement is defined as a significant positive or negative treadmill movement that is less than one second in duration. **b,** Percentage of trials with a bout of running at each bin in the trial. Running is defined as a significant negative treadmill movement (ball rolling backwards with forward mouse movement) that is at least one second in duration. **c,** Percentage of trial-time of the 3 types of movement on treadmill. Total time is 27 minutes (100 trials of 16.2 seconds - 2 baseline minutes and 14.2 seconds after onset of first odor). Statistics are paired t-tests. **d,** Number of events with amplitudes less than 10 STD per neuron per minute of recording. **e,** Number of events with amplitudes greater than 10 STD per neuron per minute. **f,** Success rate of ‘plateau-like’ events that occurred during the 3 types of movements. **g,** Firing rate increase of ‘plateau-like’ events. **h,** Same as (a) for licking in non-match trials. **i,** Same as (h) for match trials. **j-n,** Same as (c-g), but for licking on all trial types.

**Supplemental Figure 6:**
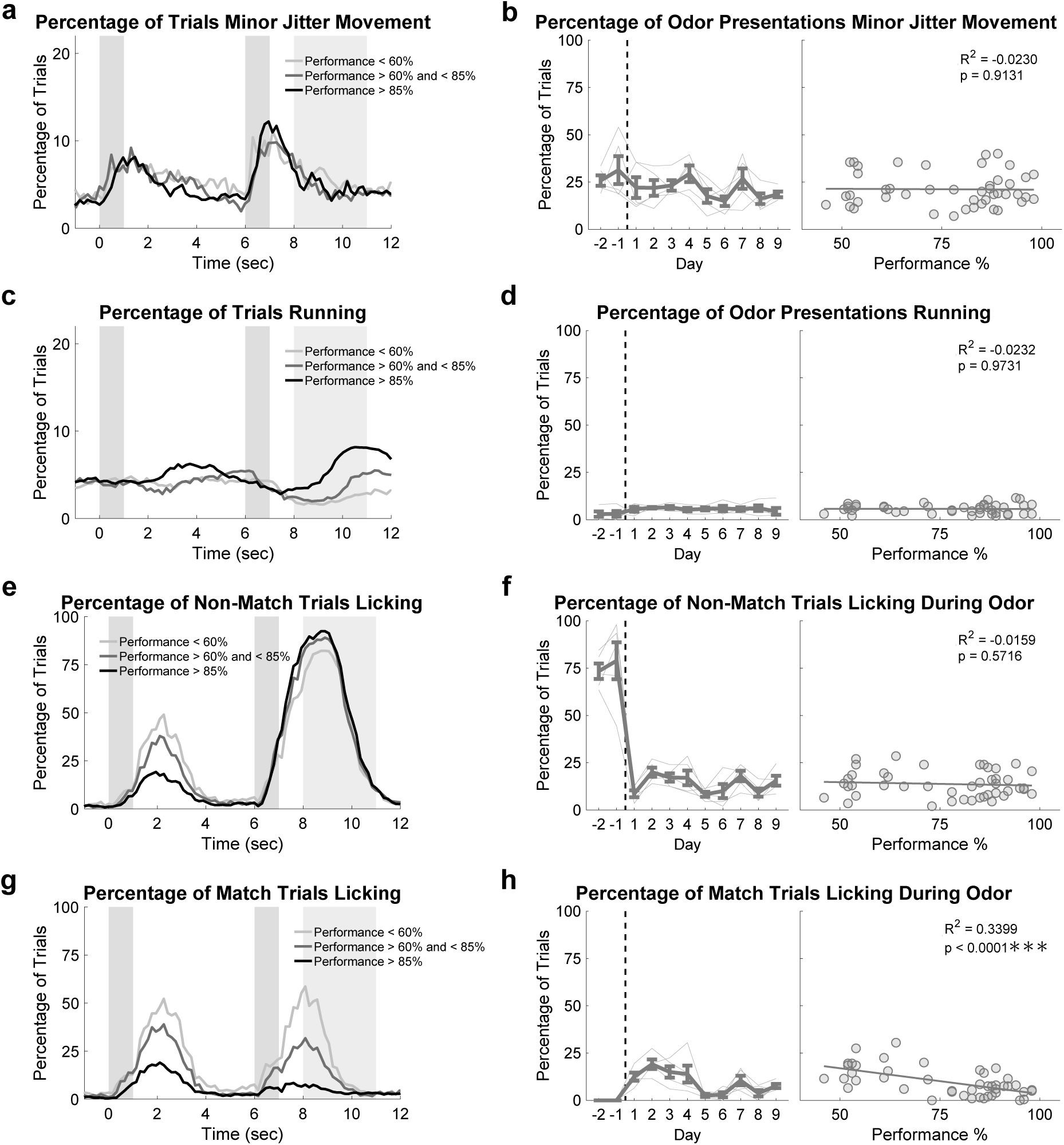
Movements do not change across learning during odor presentations. **a,** Similar to Supp. Fig. 5a, but for learning animals and split by 3 levels of performance. **b,** Percentage of odor presentations with minor jitter movements did not differ across days and performance. Statistics are the same as similar panels in Figure 3. **c-d,** Same as (a-b), but for running bouts. **e-f,** Same as (a-b), but for licking in non-match trials. **g-h,** Same as (e-f), but for match trials. The effect is related to learning to refrain from licking on these match trials, but it cannot explain any effects related to BTSP.

**Supplemental Figure 7:**
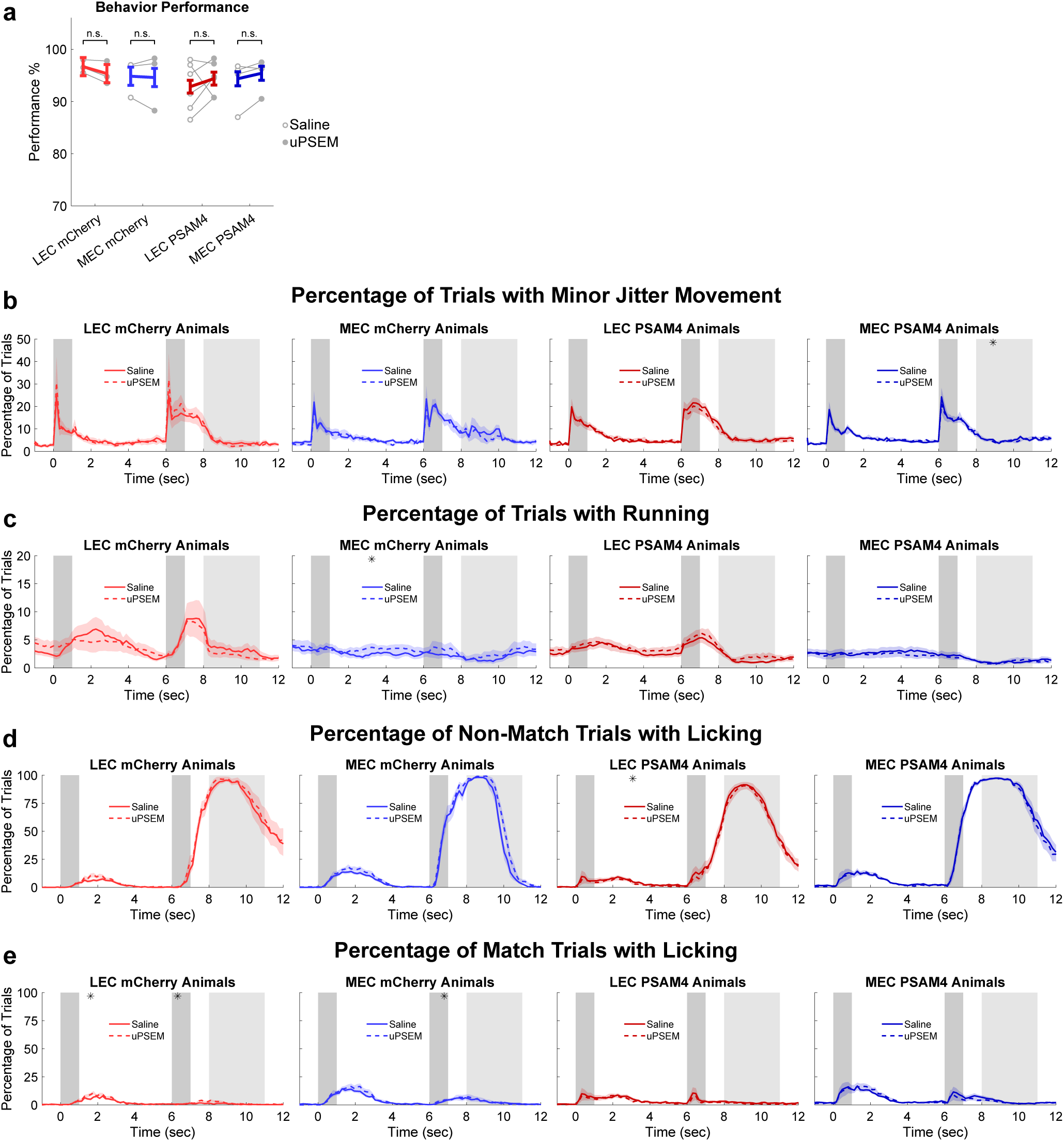
LEC or MEC inhibition had no effect on behavioral performance or movements. **a,** Average behavioral performance was unaffected by uPSEM. Statistics are the same as similar panels in Fig. 4. **b-e,** Minor jitter movements, running, and licking were largely unaffected by uPSEM. There were some minor significant bins, but these were random and cannot explain any other findings. Statistics are the same as similar panels in Fig 4.

**Supplemental Figure 8:**
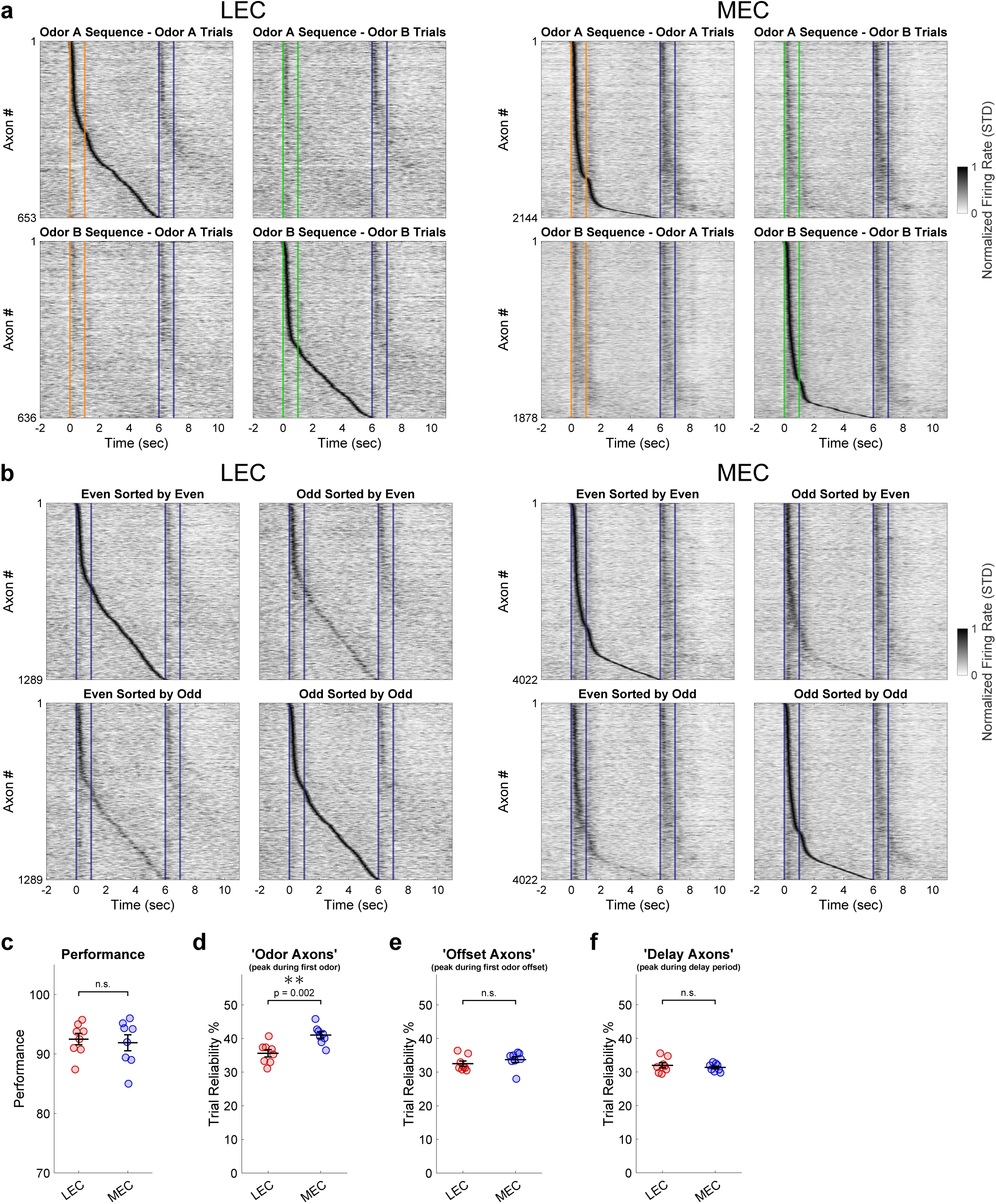
LEC and MEC sequences, and ‘odor axon’ reliability was greater for MEC. **a,** LEC and MEC sequences for expert performance split by Odor A and Odor B to show the similarities and highlight odor specificity differences. **b,** LEC and MEC preferred odor sequences for expert performance doing even/odd trial validation. The robust shadows in ‘even sorted by odd’ and ‘odd sorted by even’ panels suggest that our sequence detection algorithm performed well at identifying axons with significant fields. **c** Behavioral performance was not different between LEC and MEC axon animals. **d-f,** Same as Fig. 5h, but for trial reliability (percentage of trials with a calcium event at its field on the preferred odor trials). Statistics are the same as Fig. 5h.

